# FFPE-CUTAC: A Single Assay, Multiple Layers

**DOI:** 10.64898/2026.09.21.753320

**Authors:** Yiyang Niu, Qunzhi Xu, Chun Yin Mak, Yiling Xu, Aditya Parmar, Alex Zevin, Nadiya Khyzha, Ronald Paranal, Eric Holland, Kami Ahmad, Steven Henikoff, Ye Zheng

## Abstract

**Background:** Archival biobanks of formalin-fixed paraffin-embedded (FFPE) specimens represent a vast and underused resource for retrospective molecular studies linked to clinical follow-up. However, fixation-induced chemical modification and nucleic acid fragmentation limit scalable genomic and transcriptomic profiling. FFPE-CUTAC (Cleavage Under Targeted Accessible Chromatin) is an RNA polymerase II-targeted DNA assay specifically designed to profile regulatory activity in FFPE tissue. Here, we assess its archival robustness, information content, compatibility with RNA-seq, and ability to recover copy-number alterations.

**Results:** In meningioma and breast cancer cohorts, we benchmarked FFPE-CUTAC against matched fresh-frozen RNA-seq, FFPE RNA-seq and FFPE whole-genome sequencing. For FFPE specimens spanning up to 25 years, FFPE-CUTAC showed no systematic collection-year-associated decline in library yield or signal-quality metrics. In contrast, FFPE RNA-seq is more affected by the FFPE tissue degradation and specimen age on read composition, gene-body coverage, and gene detection. By measuring RNA polymerase II occupancy on chromatin rather than mature RNA abundance, FFPE-CUTAC is less constrained by transcript half-life, polyadenylation or transcript-capture design. FFPE-CUTAC extends beyond gene regions to non-coding regulatory elements, while the shared gene-level signals supported its integration with existing RNA-seq cohorts for large-scale, long-term clinical association study. FFPE-CUTAC libraries also retain DNA dosage information and can be used to recover chromosome-arm gain, loss and intact states with 96% concordance to matched whole-genome sequencing.

**Conclusions:** By converting routinely preserved pathology sections into integrated regulatory and copy-number profiles, FFPE-CUTAC provides a practical foundation for constructing clinically annotated disease maps to support molecular stratification, prognostic modeling and treatment-association studies.

## Background

The world’s largest tissue archive is already in our hands, but most of its molecular information remains locked in paraffin^1^. For more than a century, laboratories have preserved human and animal tissues as formalin-fixed, paraffin-embedded (FFPE) blocks^2–4^. This has created tens of billions of specimens across cancer, neurodegeneration, cardiovascular disease, infection, inflammation, transplantation, and toxicology^2–11^. Unlike prospective molecular cohorts that rely on fresh tissues, these FFPE blocks are linked to years of follow-up, including histology, diagnosis, long-term treatment responses, and survival^2, 12, 13^. Yet the field is stuck, and the FFPE resources are largely untapped, not because FFPE blocks are simply “old tissue”. It is because the same chemistry that makes FFPE compatible with routine pathology and easy storage at room temperature also fragments, crosslinks, and chemically modifies nucleic acids^14–18^.

RNA-seq and whole-genome sequencing (WGS) are common assays to profile FFPE specimens, but each faces fundamental quality barriers that goes beyond cost. FFPE RNA-seq is particularly vulnerable to fixation-induced RNA damage. Formalin-induced fragmentation, chemical modification and crosslinking reduce RNA recovery and produce uneven transcript coverage. DV200, defined as the fraction of RNA fragments longer than 200 nucleotides, is widely used to assess RNA quality and sequencing suitability. DV200 values were found to vary substantially across FFPE blocks, indicating reduction in library yield, greater coverage bias and less reliable gene-level quantification^19–21^. These limitations are amplified at low RNA input and can reduce gene detection and concordance with matched fresh-frozen (FF) RNA-seq^22, 23^. Capture-based protocols can improve transcript recovery from degraded RNA, but restrict measurement to predefined regions^22–24^. Beyond these technical limitations, conventional gene-centric RNA-seq provides limited information on regulatory activity outside annotated gene bodies^25–27^. Poly(A)-selected libraries also underrepresent non-polyadenylated transcripts^28^. More fundamentally, steady-state RNA abundance is an indirect measure of transcriptional activity because it reflects both RNA synthesis and decay. This distinction is particularly important for unstable RNA species, whose mature abundance can diverge substantially from ongoing transcription activities^29–31^.

FFPE WGS can detect mutations, structural variants and copy-number alterations (CNAs), but fixation-related DNA damage can compromise accuracy. Cytosine deamination and other formalin-induced lesions generate false-positive variants and characteristic mutational signatures that may require molecular repair or computational filtering^16–18, 32^. Uneven and GC-dependent coverage can also complicate copy-number and structural-variant inference^17, 32–34^. Deep WGS remains the most comprehensive assay for genome-wide variation^35, 36^, but its sequencing, storage and computational requirements constrain routine application to large FFPE cohorts^17, 32, 37^. Moreover, WGS does not directly measure active transcriptional or regulatory states, limiting its use as a stand-alone assay when both regulatory activity and CNAs must be obtained from limited tissue.

Routine clinical pathology combines H&E morphology with targeted molecular biomarkers, but these measurements do not provide an unbiased genome-wide view of regulatory activity and chromosomal dosage from the same tissue region^38–42^. Transcriptional regulatory programs define tumor and tumor microenvironment states, whereas CNAs and aneuploidy reflect chromosomal instability and tumor evolution^27, 43–48^. Obtaining both measurements by pairing FFPE RNA-seq with FFPE WGS requires separate workflows and substantial tissue input, cost and turnaround time^17, 22, 23^. In practice, the two assays together can consume the equivalent of eight to sixteen 5 *µ*m sections and cost approximately US$1,100–2,600 per sample (Table 1). These requirements constrain molecular profiling of limited clinical specimens and large retrospective cohorts, where tissue preservation, affordability and timely results are essential^49–53^. This gap motivates a single FFPE-compatible assay that can recover genome-wide transcriptional regulatory programs and copy-number states from minimal tissue at a practical cost.

**Table 1:** Per-sample input requirements, experimental and computational cost, profiling scope, and workflow across FF and FFPE assays.

| Per sample | FF WGS | FF RNA-seq | FFPE WGS | FFPE RNA-seq | FFPE-CUTAC |
| --- | --- | --- | --- | --- | --- |
| **Tissue volume** | 10–50 mg fresh frozen | 10–50 mg fresh frozen | 50–300 mm<sup>2</sup> × 20–40 μm | 50–300 mm<sup>2</sup> × 20–40 μm | < 25 mm<sup>2</sup> × 5 μm*<br>(multiple dissection samples per slide). |
| **Extraction** | ~\$28–45 | ~\$45–50 | ~\$38–75 | ~\$56–75 | N/A |
| **Library + sequencing** | \$750–2,180 (50–120× coverage) | ~\$150–270 | \$750–2,180 (50–120× coverage) | ~\$250–270 | ~\$50–100 |
| **Total cost/sample** | ~\$780–2,225 | ~\$195–320 | ~\$788–2,255 | ~\$306–345 | ~\$50–100 |
| **Profiling** | Whole genome | Whole transcriptome | Whole genome | Capture-defined coding transcriptome | Genome-wide targeted chromatin |
| **Workflow** | Multi-step extraction + library | Multi-step + reverse transcription | Multi-step extraction + library | Multi-step + reverse transcription | Streamlined on-slide + single-tube |
| **Mean aligned BAM storage\*\*** | ~244 GB | ~2.6 GB | ~244 GB | ~2.9 GB | ~1.2 GB |
| **Mean CNA calling runtime\*\*\*** | ~46 min | ~3.3 min | ~46 min | ~3.8 min | ~0.5 min |

\* FFPE-CUTAC protocol has been tested feasible on FFPE slides of 4-10 $\mu$ m.
\*\* Storage values are the average of aligned BAM file sizes measured in this study.
\*\*\* Runtime is the average wall-clock time starting from aligned BAM files to chromosome-arm calls using each assay-specific pipeline per-sample.

FFPE-CUTAC (Cleavage Under Targeted Accessible Chromatin)^3, 54^ provides such a framework. FFPE-CUTAC directly maps RNA polymerase II (RNAPII) occupancy genome-wide by targeting the phosphorylated C-terminal domain. The human RNAPII C-terminal domain contains 52 heptad repeats built around the YSPTSPS consensus, providing multiple Ser5-phosphorylated epitopes for antibody targeting^55^. Because this consensus lacks lysine, RNAPII is less susceptible to fixation-associated epitope masking than lysine-rich nucleosomal targets^56^. Antibody-tethered Tn5 preferentially releases approximately 120-bp DNA fragments adjacent to RNAPII, making the assay compatible with the fragmented DNA characteristics of FFPE specimens and less dependent on long intact nucleic-acid molecules^3^. RNAPII occupancy therefore provides a direct chromatin readout of engaged transcriptional machinery that is less constrained by RNA integrity, transcript half-life, polyadenylation or transcript capture. The resulting signal spans promoters, gene bodies and non-coding regulatory elements, including enhancer-associated transcription that is not fully captured by gene-centric RNA-seq^25, 26^. Genome-wide coverage of the FFPE-CUTAC libraries also retains DNA-dosage information, potentially enabling recovery of chromosome-arm aneuploidy states from the same library used for regulatory profiling. FFPE-CUTAC operates directly on a single 5 *µ*m FFPE section without nucleic-acid extraction and typically uses less than 25 mm^2^ of tissue, representing a 10-to 100-fold reduction in tissue input relative to FFPE RNA-seq or FFPE WGS (Table 1). FFPE-CUTAC therefore provides a low-input, shallow-sequencing workflow suited to limited clinical specimens and large retrospective FFPE cohorts.

In this study, we benchmark FFPE-CUTAC against matched FF RNA-seq, FFPE RNA-seq and FFPE WGS across two complementary archival tumor cohorts. We compare data quality, sensitivity to FFPE-associated degradation, genomic coverage and information content. We then identify transcriptional regulatory signals that are underrepresented by RNA-seq. At the same time, we determine whether shared gene-level signals support integration with existing RNA-seq reference cohorts linked to clinical follow-ups and molecular annotations^57, 58^. Finally, we test whether FFPE-CUTAC readout recovers WGS-defined CNA states. Together, these analyses evaluate whether a single low-input FFPE-CUTAC assay can provide practical, robust and multi-layer molecular profiling from routine archival specimens.

## Results

### FFPE-CUTAC robustly captures broad transcriptional regulatory information from FFPE tissue

We used two FFPE tumor cohorts to benchmark FFPE-CUTAC against RNA-seq and WGS across complementary tissue settings^54, 58^ (Fig. 1A). The meningioma cohort included 30 patients with matched FFPE-CUTAC, FFPE RNA-seq, and FFPE WGS data. A subset of 17 patients also had matched fresh-frozen poly(A)-selected RNA-seq data. Most meningiomas were histologically benign and represented a relatively circumscribed tumor setting with generally high tumor purity. These specimens were collected between 2017 and 2023. The breast cancer cohort included 15 patients with FFPE RNA-seq, of whom 13 were also profiled by FFPE-CUTAC. The specimens were collected between 1999 and 2012 and included HER2^+^, ER^+^, and triple-negative breast cancer (TNBC) subtypes. This cohort represented an older and more heterogeneous invasive tumor setting, with variable tumor purity due to stromal and immune admixture. Together, these cohorts enabled direct cross-modality benchmarking and assessment of FFPE-CUTAC performance across broad tissue collect period with contrasting tissue compositions.

**Figure 1.**
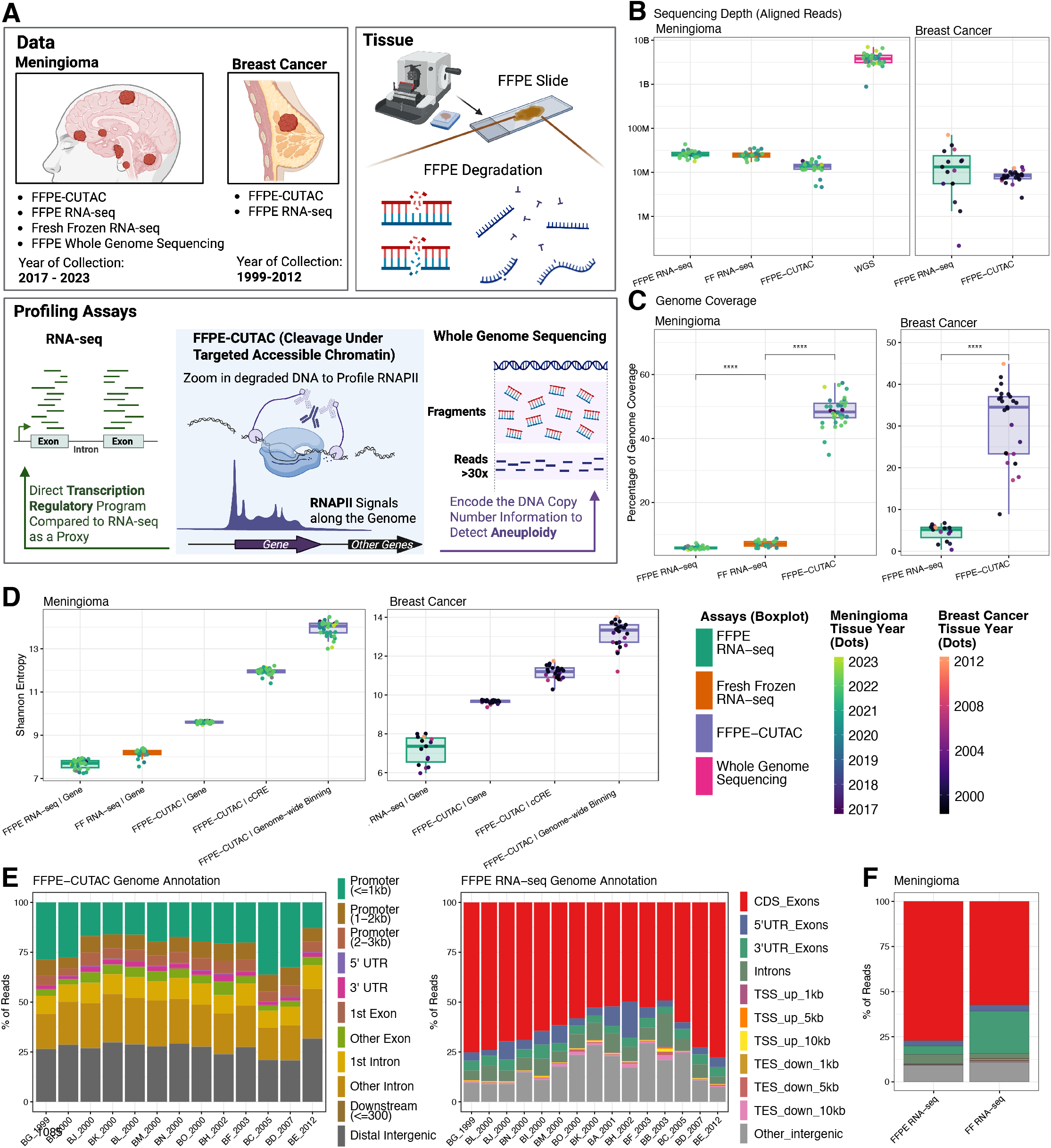
Cross-assay benchmarking of data quality and genomic information coverage. **A.** Study overview showing the meningioma and breast cancer cohorts, tissue collection periods, FFPE-associated nucleic acid degradation, and assay-specific readouts. The meningioma cohort included FFPE-CUTAC, FFPE RNA-seq, fresh frozen (FF) RNA-seq, and FFPE whole-genome sequencing (WGS). The breast cancer cohort included FFPE-CUTAC and FFPE RNA-seq. **B.** Number of aligned reads per library by assay and cohort, displayed on a log_10_ scale. **C.** Percentage of the genome covered after downsampling each RNA-seq and FFPE-CUTAC library to 5 million aligned reads, shown by assay and cohort. **D.** Shannon entropy^59^ calculated from raw counts across genes for RNA-seq and across genes, candidate cis-regulatory elements (cCREs)^26^, or consecutive 500-bp genome-wide bins for FFPE-CUTAC. Meningioma results include FFPE and FF RNA-seq, whereas breast cancer results include FFPE RNA-seq. **E.** Genomic-feature annotation of aligned reads from breast cancer FFPE-CUTAC and FFPE RNA-seq libraries. FFPE-CUTAC reads were assigned to ChIPseeker^119^ categories after biological replicates were merged within each patient. FFPE RNA-seq reads were assigned using RSeQC^120^. Each stacked bar represents one patient, ordered by tissue collection year. **F.** Cohort-level mean RSeQC read-distribution profiles for meningioma FFPE RNA-seq and FF RNA-seq libraries. Point colors in **B–D** indicate tissue collection year. In box plots, center lines denote medians, box limits denote the first and third quartiles, and whiskers extend to the most extreme values within 1.5 times the interquartile range. Points represent individual libraries or samples, as applicable. Brackets denote two-sided Wilcoxon rank-sum tests for the indicated pairwise comparisons. * *P <* 0.05, ** *P <* 0.01, *** *P <* 0.001, and **** *P <* 0.0001.

We started by comparing the basic data characteristics across modalities. As expected, WGS libraries had the highest sequencing depth, while FFPE-CUTAC had even shallower libraries than matched RNA-seq assays (Fig. 1B). After downsampling each library to five million aligned fragments, FFPE-CUTAC covered a substantially larger fraction of the mappable genome than RNA-seq (Fig. 1C). This difference reflected assay scope. RNA-seq reads were concentrated within annotated transcripts, whereas FFPE-CUTAC captured RNAPII-associated DNA fragments across promoters, gene bodies, and non-coding regulatory regions. We next calculated Shannon entropy^59^ from raw feature counts to quantify the information content across each defined feature space (Fig. 1D). When signal was summarized over the same annotated genes, FF RNA-seq showed higher entropy than FFPE RNA-seq, consistent with the expected loss of transcriptome quality in formalin-fixed tissue. FFPE-CUTAC showed higher gene-level entropy than either RNA-seq assay. Within FFPE-CUTAC, entropy increased as the feature space expanded from genes to candidate cis-regulatory elements (cCREs)^26^ and genome-wide 500-bp bins. Thus, although FFPE-CUTAC used fewer reads, its DNA-based regulatory footprint covers a broader feature space and more information than RNA-seq.

We then asked whether archival age altered the genomic distribution of reads. Among breast cancer specimens, which are relatively older, FFPE-CUTAC showed broadly stable genomic-feature composition across collection years (Fig. 1E). This stability was also evident in tissue-age group summaries for breast cancer and per-sample annotation profiles for meningioma (Supplementary Fig. 1A-B). In contrast, FFPE RNA-seq showed collection-year-associated redistribution across genomic features. The coding sequence (CDS) exon representation decreased and rebounded across the archival series, accompanied by an inverse pattern in the intergenic read fraction (Fig. 1E and Supplementary Fig. 1C-D). Matched meningioma data further showed distinct read-distribution profiles between FF poly(A)-selected RNA-seq and FFPE RNA-exome capture assay (Fig. 1F). FF libraries had lower CDS-exon representation and higher 3*^′^* untranslated region (UTR) exon representation than FFPE libraries. Variation in CDS-exon, UTR, intronic, and intergenic read fractions likely reflects the combined effects of specimen preservation and library-enrichment strategy.

We therefore further examined gene-body coverage from the 5*^′^* end to the 3*^′^* end (Fig. 2A). FF meningioma RNA-seq showed a more uniform profile across gene bodies. In contrast, FFPE RNA-seq from both meningioma and breast cancer showed a 5*^′^*-skewed coverage. The more recently collected meningioma FFPE libraries showed less between-sample variation (light green thin lines closer to the average trend in Fig. 2A) than the older breast cancer FFPE libraries (dark thin lines in Fig. 2A), showing the FFPE RNA-seq sensitivity to specimen age and degradation. We then quantified coverage uniformity using the transcript integrity number (TIN), which summarizes the evenness of mapped-read coverage across each transcript^60^. Median TIN was lower in FFPE RNA-seq libraries than in matched FF RNA-seq libraries at both the sample and transcript levels (Fig. 2B and Supplementary Fig. 1E). TIN declined progressively with transcript length in FFPE libraries (Fig. 2C and Supplementary Fig. 1F). In FF libraries, however, TIN remained relatively stable across short and intermediate transcripts and declined mainly among the longest transcripts due to technical challenges of reverse transcribing long RNA. The consistent dropping patterns in FFPE RNA libraries indicate greater length-dependent coverage non-uniformity. Among the transcript properties evaluated, 3*^′^* UTR length showed the strongest association with paired TIN loss between matched FFPE and FF meningioma libraries (Supplementary Fig. 1G-H). RNA half-life provided additional stratification. In FFPE libraries of breast cancer, transcripts with shorter reference half-lives had lower TIN, and TIN increased with more recent collection year across half-life groups (Fig. 2D). A similar half-life ordering was observed in meningioma, where FF libraries had higher TIN than matched FFPE libraries across all half-life groups (Supplementary Fig. 1I).

**Figure 2.**
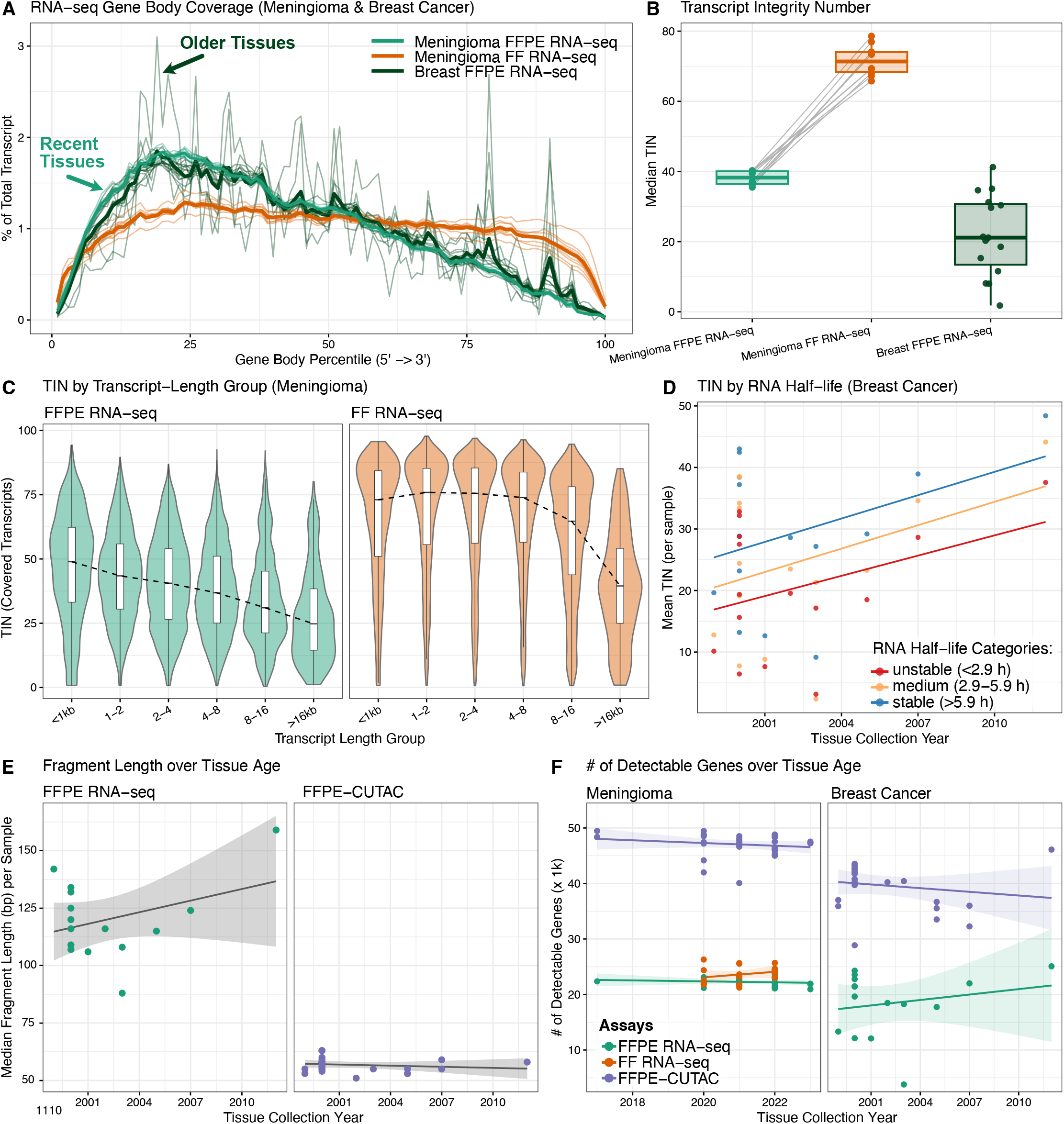
RNA integrity, fragment length, and gene detection across archival tumor cohorts. **A.** 5*^′^*-to-3*^′^* gene-body coverage profiles^120^ for meningioma FFPE RNA-seq, meningioma fresh frozen (FF) RNA-seq, and breast cancer FFPE RNA-seq. Thin lines represent individual libraries, and bold lines represent group means. **B.** Median transcript integrity number (TIN) per RNA-seq library. Matched meningioma FFPE and FF RNA-seq libraries are connected by gray lines. Breast cancer FFPE RNA-seq libraries are shown separately. **C.** TIN distributions of covered transcripts stratified by transcript-length group for meningioma FFPE and FF RNA-seq libraries. Violin plots show transcript-level distributions, internal box plots summarize each group, and dashed lines connect group medians. **D.** Mean TIN per breast cancer FFPE RNA-seq library for genes grouped by RNA half-life from RNAdecayCafe^100^. Genes were classified as unstable (*<* 2.9 h), medium (2.9–5.9 h), or stable (*>* 5.9 h). **E.** Median fragment length per breast cancer FFPE RNA-seq and FFPE-CUTAC library as a function of tissue collection year. **F.** Number of detectable genes per library, as a function of tissue collection year, shown by assay for the meningioma and breast cancer cohorts. A detectable gene was defined as having at least one assigned read or fragment. In box plots, center lines denote medians, box limits denote the first and third quartiles, and whiskers extend to the most extreme values within 1.5 times the interquartile range. Points represent individual libraries. In **D–F**, fitted lines are ordinary least-squares regressions, with shaded bands, where shown, indicating 95% confidence intervals.

We then compared fragment lengths across assays. In FFPE RNA-seq libraries of breast cancer, median insert length increased with collection year, consistent with greater RNA fragmentation in older specimens (Fig. 2E). In contrast, median FFPE-CUTAC fragment length varied within a narrow range and showed no clear association with collection year (Supplementary Fig. 2A-E). The FFPE-CUTAC pattern is consistent with pA–Tn5 tagmentation, which releases short DNA fragments adjacent to RNAPII-occupied loci (*∼*120bp) and reduces dependence on long, intact nucleic-acid molecules. We next assessed whether collection year was associated with gene-level signal recovery (Fig. 2F). In the breast cancer cohort, FFPE RNA-seq libraries showed a slightly increasing pattern in the number of detected genes over collection years. FFPE-CUTAC remained comparatively more stable and showed larger amounts of gene detection in both cohorts. Similarly, at the peak regions level, FFPE-CUTAC peak numbers and fractions of reads in peaks (FRiP) showed no systematic decline with specimen age for both meningioma and breast cancer samples (Supplementary Fig. 2F-I). Finally, GC-content distributions showed no evident systematic shift with collection year within FFPE-CUTAC, RNA-seq, or WGS libraries (Supplementary Fig. 2J-K). RNA-seq and FFPE-CUTAC reads were shifted toward higher GC content than WGS reads, consistent with their enrichment within transcribed and promoter-proximal genomic regions. Thus, the stable FFPE-CUTAC profiles were not accompanied by an age-associated shift in library GC composition.

Together, these analyses establish that FFPE-CUTAC retains a broad RNAPII signal across recent and older FFPE specimens. FFPE RNA-seq libraries showed collection-year-associated declines in read composition, transcript coverage uniformity, insert length, and gene detection. In contrast, FFPE-CUTAC required less input tissue material and covered a broader fraction of the genome at even shallower sequencing depth and showed no systematic age-associated decline in genomic-feature composition, fragment length, gene-associated signal, or peak-level metrics. These findings support FFPE-CUTAC as a robust assay for regulatory profiling of archival FFPE tissue.

### FFPE-CUTAC uniquely profiles the regulatory activities that are ‘under-represented by RNA-seq

To compare FFPE-CUTAC and RNA-seq on a common genomic unit, we quantified both assays over annotated genes (Fig. 3A and Supplementary Fig. 3A). This represents a conservative comparison for FFPE-CUTAC because it excludes signal at non-coding regulatory elements. In matched meningioma samples, FFPE-CUTAC and FF RNA-seq identified a large set of co-detected genes and a distinct group with high FFPE-CUTAC readout but low RNA-seq signal (Fig. 3A). Comparable patterns were observed when FFPE-CUTAC was compared with meningioma and breast cancer FFPE RNA-seq (Supplementary Fig. 3B–E). We defined genes with high FFPE-CUTAC and low RNA-seq counts as FFPE-CUTAC-favored genes and examined their GENCODE biotypes^61^. These genes spanned multiple biotypes, with protein-coding genes forming the largest component, followed by antisense and lincRNA (i.e., long, intervening noncoding RNA) genes (Fig. 3B, C). Biotype-stratified signal distributions showed similar patterns across the meningioma and breast cancer comparisons (Supplementary Fig. 3F–H and Supplementary Fig. 4A–D).

**Figure 3.**
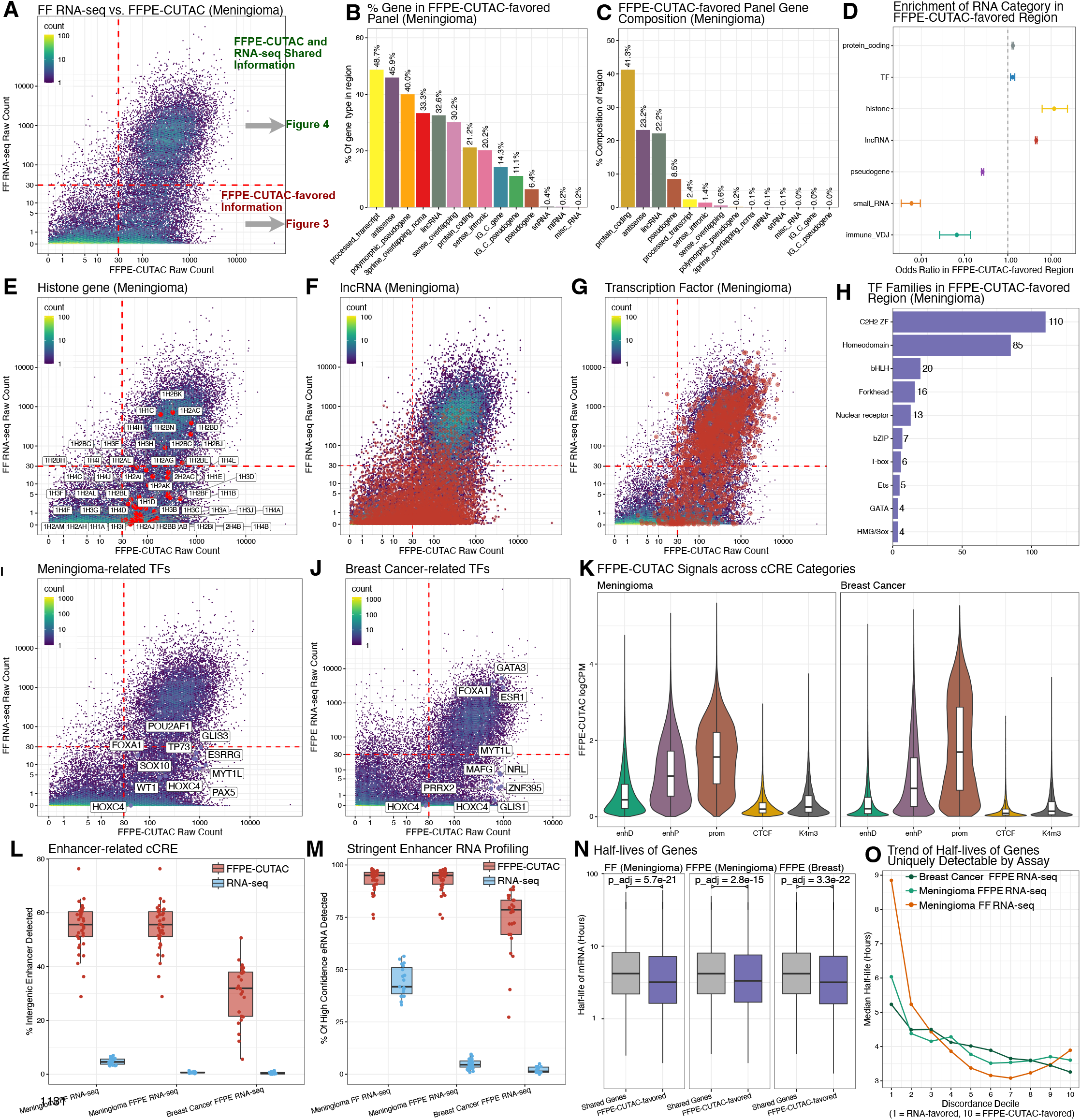
Gene and regulatory features detection by FFPE-CUTAC relative to RNA-seq. **A.** Mean raw count per gene across matched meningioma FFPE-CUTAC and FF RNA-seq libraries. Red dashed lines mark the 30-count thresholds. Genes with a mean FFPE-CUTAC count *>* 30 and a mean FF RNA-seq count *<* 30 were classified as FFPE-CUTAC-favored. Genes with mean counts *>* 30 in both assays were classified as shared. **B.** Percentage of genes within each GENCODE biotype^61^ classified as FFPE-CUTAC-favored. **C.** GENCODE biotype composition of the FFPE-CUTAC-favored gene set. **D.** Enrichment of broad gene categories in the FFPE-CUTAC-favored region using logCPM-normalized counts. Points indicate odds ratios from Fisher’s exact tests comparing FFPE-CUTAC-favored genes with shared genes. Horizontal bars indicate 95% confidence intervals. The vertical dashed line marks an odds ratio of 1. **E-G.** Gene-level count-density plots with histone genes (**E**), long non-coding RNA (lncRNA) genes (**F**), and transcription factor (TF) genes^77^ (**G**) highlighted in red. Common HIST-family prefixes were omitted from histone gene labels for readability. **H.** Structural family distribution of TF genes in the meningioma FFPE-CUTAC-favored region. Bar-end labels indicate the number of FFPE-CUTAC-favored TF genes in each family. **I.** Selected meningioma-associated TFs labeled on the meningioma FFPE-CUTAC and FF RNA-seq comparison. **J.** Selected breast cancer-associated TFs labeled on the breast cancer FFPE-CUTAC and FFPE RNA-seq comparison. **K.** Distribution of FFPE-CUTAC logCPM values among different categories of cCREs regions^26^ and cohorts. cCRE classes include distal enhancer-like (enhD), proximal enhancer-like (enhP), promoter-like (prom), CTCF-only (CTCF), and H3K4me3-only (K4m3) cCREs. **L.** Percentage of intergenic enhancer-like cCREs detected per library by FFPE-CUTAC and RNA-seq in the indicated meningioma and breast cancer comparisons. **M.** Percentage of high-confidence enhancer loci detected per library for the same assay and cohort comparisons as in **L**. High-confidence loci were defined as intergenic enhD or enhP cCREs overlapping FANTOM5 permissive enhancers^121^, excluding readthrough candidates and requiring bidirectional enhancer RNA evidence (Methods). **N.** mRNA half-life^100^, displayed on a log_10_ scale, for shared and FFPE-CUTAC-favored genes. Comparisons are shown for meningioma FF RNA-seq, meningioma FFPE RNA-seq, and breast cancer FFPE RNA-seq. Brackets indicate Benjamini-Hochberg-adjusted *P* values from two-sided Wilcoxon rank-sum tests. **O.** Median mRNA half-life across deciles of FFPE-CUTAC-versus-RNA-seq discordance. Decile 1 represents RNA-seq-favored genes, and decile 10 represents FFPE-CUTAC-favored genes. In box plots, center lines denote medians, box limits denote the first and third quartiles, and whiskers extend to the most extreme values within 1.5 times the interquartile range. Points represent individual libraries where shown. Violin widths represent kernel-density estimates and contain embedded box plots.

We next grouped genes into broad RNA categories and tested their abundance among FFPE-CUTAC-favored genes. Histone genes showed the strongest enrichment, followed by lncRNAs and transcription factors (TFs) (Fig. 3D and Supplementary Fig. 4E-F). Replication-coupled histone genes provided the clearest mechanistic example. In matched meningioma samples, many histone genes showed strong FFPE-CUTAC signal but weak FF RNA-seq signal (Fig. 3E). These genes are transcribed by RNAPII^62^, but their mature mRNAs generally lack canonical poly(A) tails^28, 63^. Instead, they undergo specialized 3*^′^*-end processing^64, 65^. FFPE-CUTAC therefore captures active RNAPII occupancy at histone loci that are underrepresented by poly(A)-selected RNA-seq. This is biologically important because histone production is coupled to DNA replication, and RNAPII at S-phase-dependent histone genes has been associated with tumor grade and recurrence in meningioma and with breast cancer proliferation-related states^54, 66^. lncRNA loci also showed stronger FFPE-CUTAC than RNA-seq signal (Fig. 3F). Some lncRNAs, such as MALAT1 and TERC, lack canonical poly(A) tails or exist in both polyadenylated and non-polyadenylated forms^67, 68^ and may therefore be underrepresented by oligo-dT-based FF RNA-seq. More broadly, lncRNAs are often low abundance^61, 69^, incompletely processed^69–71^, or retained in the nucleus and chromatin compartments^61, 72–74, 74, 75^, which can reduce their representation in steady-state RNA-seq. Selected nuclear-retained and chromatin-associated lncRNAs, including XIST, NEAT1, MALAT1, AIRN, KCNQ1OT1, TERC, and TUG1, illustrated this pattern across the meningioma and breast cancer comparisons (Supplementary Fig. 4G-I). FFPE-CUTAC does not measure mature lncRNA abundance. Instead, it measures RNAPII-associated activity at the underlying locus. It can therefore detect active lncRNA loci when steady-state RNA signal is weak. For comparisons with FFPE RNA-seq, capture-panel coverage may also contribute to reduced lncRNA representation.

Moreover, TF genes were also enriched among FFPE-CUTAC-favored loci and represented major DNA-binding families, including C2H2 zinc-finger, homeodomain, bHLH, fork-head, nuclear receptor, ETS, GATA, and HMG/SOX families^76, 77^ (Fig. 3G-H and Supplementary Fig 4J-K). RNAPII occupancy at these loci provides a complementary measure of TF-gene activity when steady-state RNA abundance is low. In meningioma, FFPE-CUTAC-favored TFs included SOX10, WT1, FOXA1, PAX5, and MYT1L (Fig. 3I), consistent with lineage and enhancer programs previously implicated in meningioma^78–82^. These signals may reflect tumor-intrinsic programs or cellular components of the tumor microenvironment. For example, MYT1L is linked to neuronal identity maintenance, and PAX5 is a canonical B-cell lineage regulator^83, 84^. In breast cancer, ESR1, FOXA1, and GATA3 were detected by both FFPE-CUTAC and FFPE RNA-seq when strongly active, consistent with the ER^+^ luminal regulatory network^85–88^ (Fig. 3J). FOXA1 determines ER-chromatin interactions and chromatin accessibility, while GATA3 shapes enhancer accessibility upstream of FOXA1 and ESR1-mediated transcription^86, 87^. The additional FFPE-CUTAC-favored breast TFs were enriched for developmental and plasticity-associated families, including homeodomain, C2H2 zinc-finger, bHLH, bZIP, nuclear receptor, forkhead, HMG/SOX and ETS families (Supplementary Fig 4L). These TF classes have documented roles in developmental identity, stem/progenitor state, epithelial–mesenchymal plasticity, endocrine resistance, invasion and metastasis^89–97^. In summary, FFPE-CUTAC captures both canonical lineage programs and RNAPII activity at regulatory loci underrepresented by RNA-seq.

We next extended the comparison beyond annotated genes. FFPE-CUTAC detected RNAPII-associated signal across ENCODE cCRE classes, including distal enhancer-like (enhD), proximal enhancer-like (enhP), promoter-like, CTCF-only, and DNase–H3K4me3 elements^26^ (Fig. 3K). We focused our analysis on intergenic enhD and enhP cCREs to exclude annotated gene bodies. The quantified RNA-seq reads over the same intervals are considered candidate enhancer RNA (eRNA) signal. FFPE-CUTAC detected a greater fraction of these intergenic enhancer loci than RNA-seq for both cohort and assay comparisons (Fig. 3L and Supplementary Fig. 4M–P). We then defined a stringent set of high-confidence transcribed enhancer loci by excluding likely readthrough from nearby expressed genes (Methods). It required overlap with FANTOM5 CAGE-supported enhancers^25, 98^ and required bidirectional RNA evidence^25^ (Supplementary Fig. 4Q). FANTOM5 CAGE data support this strategy because bidirectional capped RNAs are a hallmark of active enhancers^99^. FFPE-CUTAC still detected RNAPII-associated signal at a greater fraction of these loci than RNA-seq (Fig. 3M). The results are consistent with the transient, low-abundance, and often non-polyadenylated nature of eRNAs, which limits recovery by poly(A)-selected FF RNA-seq, and with the gene-targeted design of FFPE RNA-seq.

Finally, we tested whether transcript stability contributed to assay discordance. Across comparisons with meningioma FF RNA-seq, meningioma FFPE RNA-seq, and breast cancer FFPE RNA-seq, FFPE-CUTAC-favored genes had shorter reference mRNA half-lives than shared genes^100^ (Fig. 3N). When discordance was treated as a continuous variable, median mRNA half-life decreased from RNA-seq-favored genes to FFPE-CUTAC-favored genes across deciles (Fig. 3O). These results are consistent with FFPE-CUTAC being less constrained by mature RNA stability than steady-state RNA-seq.

### Shared gene-level signal enables FFPE-CUTAC and RNA-seq integration

Building on the FFPE-CUTAC-favored signal identified above, we continued to ask whether the gene-level information shared between FFPE-CUTAC and RNA-seq could support integration with existing RNA-seq cohorts linked to disease annotations and clinical outcomes. Such integration would allow newly profiled archival FFPE specimens to leverage established transcriptomic reference maps. We therefore restricted both assays to shared annotated genes (Fig. 3A), creating a common feature space for cross-modality integration. We then tested whether RNAPII-associated FFPE-CUTAC profiles aligned with the RNA-seq-defined disease structure map and preserved clinically relevant molecular patterns.

We first jointly embedded FFPE-CUTAC, FF RNA-seq, and FFPE RNA-seq libraries within a combined meningioma reference cohort. FFPE-CUTAC profiles occupied the same broad manifold as FF RNA-seq, whereas FFPE RNA-seq libraries formed a smaller, more separated cluster (Fig. 4A-C). Study-of-origin annotations confirmed no batch effect after integration (Supplementary Fig. 5A). We next restricted the integration to FFPE-CUTAC and FF RNA-seq. Matched libraries from the same patients occupied nearly co-localization positions (Fig. 4D). The integrated space also retained established meningioma clinical grading structure (Fig. 4E). World Health Organization (WHO) grade 2 and 3 tumors were enriched in the central region, whereas grade 1 tumors were more frequent in the lower-left and right regions. Because chromosome 22q loss is one of the most common genomic alterations in sporadic meningioma and has been associated with recurrence in subsets of grade 1 disease, we examined its distribution across the embedding^101–104^ (Fig. 4F). Chromosome 22q loss was enriched in the central and right regions, whereas 22q-intact tumors were more frequent in the lower-left region. NF2 is located on chromosome 22q and encodes Merlin, a tumor suppressor that defines a major molecular axis of meningioma biology^101, 102, 105^. Consistent with the chromosome 22q pattern, the NF2/Merlin signature was higher in the predominantly 22q-intact lower-left region and lower in the 22q-loss-enriched central and right regions (Fig. 4G). DNA methylation groups provided an independent molecular annotation^105^. Hypermitotic, immune-enriched, and Merlin-intact tumors were enriched in the central, right, and left regions, respectively (Fig. 4H). These patterns were consistent with established methylation-defined meningioma states and their distinct biological and clinical properties^105–107^. Merlin-intact meningiomas have the most favorable clinical outcomes, immune-enriched meningiomas show immune infiltration, and hypermitotic meningiomas show cell-cycle-associated biology and the least favorable survival outcomes^105, 107^. The enriched, suppressed, and ratio scores derived from a 34-gene meningioma panel also showed gradients across the embedding (Supplementary Fig. 5B-D)^108^. Together, these results show that FFPE-CUTAC can be integrated with FF RNA-seq while retaining patient-level correspondence and disease-relevant meningioma molecular structure.

**Figure 4.**
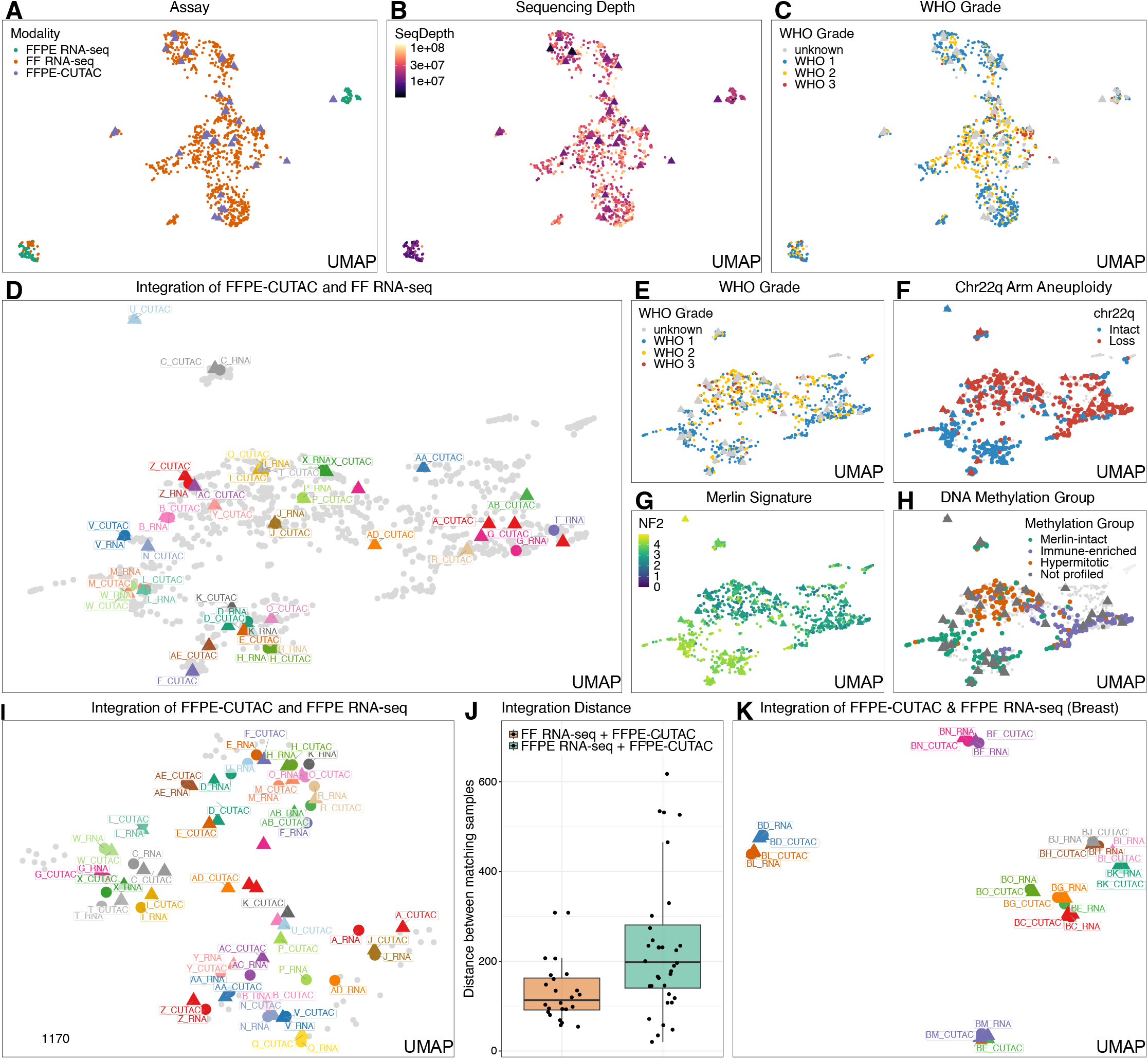
Cross-assay integration of FFPE-CUTAC and RNA-seq libraries. **A-C.** Shared UMAP embedding of meningioma fresh frozen (FF) RNA-seq, FFPE RNA-seq, and FFPE-CUTAC libraries, colored by assay (**A**), sequencing depth (**B**), and WHO grade (**C**). **D.** Shared UMAP embedding of meningioma FF RNA-seq and FFPE-CUTAC libraries, with matched patient pairs highlighted and labeled. **E-H.** Same embedding as in **D**, colored by WHO grade (**E**), chromosome 22q arm status classified as intact or loss (**F**), continuous NF2/Merlin signature score (**G**), and DNA methylation group^105^ (**H**). **I.** Shared UMAP embedding of meningioma FFPE RNA-seq and FFPE-CUTAC libraries, with matched patient pairs highlighted and labeled. **J.** Euclidean distances between matched FF RNA-seq and FFPE-CUTAC libraries (orange) and between matched FFPE RNA-seq and FFPE-CUTAC libraries (green) in the corresponding integrated embedding spaces shown in **D** and **I**. **K.** Shared UMAP embedding of breast cancer FFPE RNA-seq and FFPE-CUTAC libraries, with matched sample pairs highlighted and labeled. In box plots, center lines denote medians, box limits denote the first and third quartiles, whiskers extend to the most extreme values within 1.5 times the interquartile range, and overlaid points represent individual matched pairs.

We next tested whether FFPE RNA-seq could support integration when FF RNA-seq was unavailable. In meningioma, matched FFPE-CUTAC and FFPE RNA-seq libraries were jointly embedded, with patient-matched pairs occupying nearby positions (Fig. 4I). However, matched-pair distances were greater and more variable than in the FF RNA-seq–FFPE-CUTAC integration (Fig. 4J). Therefore, FFPE RNA-seq supports cross-modality alignment, but FF RNA-seq provides a more stable reference map for integrating FFPE-CUTAC. We applied the same framework to breast cancer. Matched FFPE-CUTAC and FFPE RNA-seq libraries aligned within a shared embedding (Fig. 4K) that separates receptor-subtype structure among ER^+^/PR*^−^*/HER2*^−^*, HER2^+^, and TNBC tumors (Supplementary Fig. 5E). No clear organization by collection year was observed (Supplementary Fig. 5F). These results support that cross-modality integration extends to older, heterogeneous FFPE tumor cohorts, but without fresh frozen samples is limited in application.

### FFPE-CUTAC data can recover chromosome-arm copy-number alterations

CNAs and aneuploidy are hallmarks of cancer genomes and are linked to tumor evolution and clinical behaviour^47, 48^. WGS remains necessary for focal mutation discovery, allele-specific copy number and complex structural variation. However, if the clinical or biological endpoint is chromosome-arm gain, loss or intact status, a shallow DNA-based RNAPII profiling assay may recover sufficient copy-number information from the same library used for transcriptional regulatory analysis. This possibility would reduce the need for separate WGS in FFPE studies focused on arm-level aneuploidy.

Therefore, we asked how well FFPE-CUTAC genome-wide coverage can be used to infer CNAs. Matched FFPE WGS served as the reference standard for chromosome-arm aneuploidy^109^. To design a CNA caller for FFPE-CUTAC, we adapted principles from established WGS and chromatin-accessibility CNA methods^27, 109–116^ (Fig. 5A). The workflow bins genome-wide FFPE-CUTAC signals to alleviate the signal distribution sparsity. It then filters unreliable genomic regions, corrects GC-content and mappability biases, smooths and segments bin-level log_2_-ratio profiles, and classifies chromosome arms as intact, gain or loss. RNA-seq is also used in practice to infer large-scale CNAs from transcriptional profiles. Thus, we compared with the CNA calls from FF RNA-seq and FFPE RNA-seq with CaSpER^117^, which infers large-scale CNAs from gene-expression and B-allele-frequency signals.

**Figure 5.**
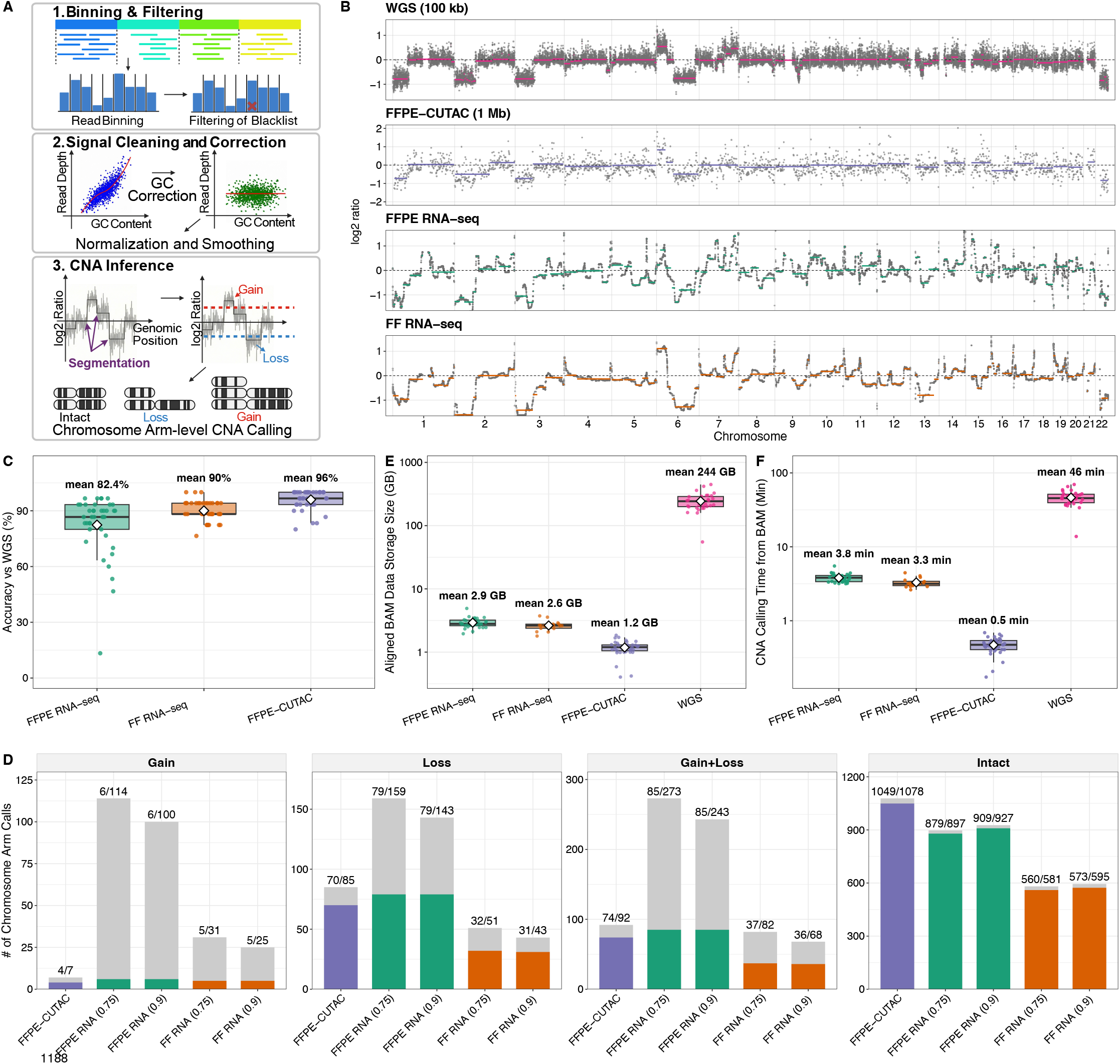
Benchmarking chromosome-arm copy-number inference across assays in meningioma. **A.** Schematic of the FFPE-CUTAC copy-number alterations detection workflow, including genomic binning and filtering, GC-content correction, normalization and smoothing, segmentation of log_2_ signal-to-baseline ratios, and chromosome-arm classification as intact, gain or loss. **B.** Genome-wide copy-number profiles for a representative meningioma patient. Tracks show FFPE WGS at 100-kb resolution as the reference^109^, FFPE-CUTAC at 1-Mb resolution, and FFPE and FF RNA-seq profiles inferred using CaSpER^117^. The x-axis spans chromosomes 1–22, and the y-axis shows log_2_ signal-to-baseline ratio. Gray points represent bin-level signals, and colored lines represent segmented profiles. **C.** Chromosome-arm concor-dance with WGS for FFPE RNA-seq, FF RNA-seq, and FFPE-CUTAC. Each point represents one of 39 chromosome arms and indicates the percentage of patients with an assay-derived CNA calls concordant with WGS. The analysis included 30 patients for FFPE-CUTAC and FFPE RNA-seq and 17 patients for FF RNA-seq. White diamonds and labels indicate mean concordance across chromosome arms. **D.** Chromosome-arm calls stratified by gain, loss, combined gain or loss, and intact status. Gray bars indicate the total number of calls assigned to each category, and colored bars indicate the subset concordant with WGS. Bar labels show the number of concordant calls over the total number of calls. **E.** Aligned BAM file size per library for each assay, shown on a log_10_ scale. White diamonds and labels indicate mean storage size. **F.** Processing time from aligned BAM files to chromosome-arm copy-number calls for each assay, shown on a log_10_ scale. White diamonds and labels indicate mean processing time. For box plots, center lines denote medians, box limits denote the first and third quartiles, and whiskers extend to the most extreme values within 1.5 times the interquartile range. Points represent individual libraries or samples, as applicable.

In a representative meningioma sample, FFPE-CUTAC read depth at 1-Mb resolution closely recapitulated the broad copy-number profile observed by matched WGS at 100-kb resolution, particularly at chromosome-arm and large-segment scales (Fig. 5B). Similar agreement between FFPE-CUTAC and WGS was observed across the cohort (Supplementary Figs. 6–7). Across 39 autosomal chromosome arms, the average aneuploidy concordance with WGS was 96.0% for FFPE-CUTAC, compared with 90.0% for FF RNA-seq and 82.4% for FFPE RNA-seq (Fig. 5C). Per-patient and per-arm analyses confirmed consistently high FFPE-CUTAC concordance with WGS across the cohort (Supplementary Fig. 8A-B). State-specific analysis showed that RNA-seq-based inference produced substantially more gain and loss calls, but only a small proportion agreed with WGS (Fig. 5D and Supplementary Fig. 8C). This discrepancy was most pronounced for FFPE RNA-seq. Thus, the FFPE-CUTAC library can provide accurate and stable aneuploidy detection.

Genomic bin size could potentially affect the FFPE-CUTAC copy-number calling performance. Across bin sizes from 100 kb to 5 Mb, we found 500-kb and 1-Mb bins yielded the highest concordance with WGS, at 95.9% and 96.0%, respectively (Supplementary Fig. 9A). Recovery of WGS-defined gains and losses was also strongest at these intermediate resolutions (Supplementary Fig. 9B). At 1-Mb resolution, replicate libraries showed a mean chromosome-arm consistency of 94.1% (Supplementary Fig. 9C). Representative profiles illustrated the expected trade-off (Supplementary Fig. 9D-E). Smaller bins increased local noise, whereas larger bins reduced genomic resolution. We therefore selected 1 Mb as the primary resolution for chromosome-arm CNA calling based on its concordance, signal stability, and reproducibility.

Beyond the direct costs associated with deeper sequencing, WGS also imposed substantial downstream data storage and computational requirements. These less visible resource demands become increasingly consequential when profiling large retrospective cohorts. Mean aligned-BAM size was 1.2 GB for FFPE-CUTAC and 244 GB for WGS, representing an approximately 200-fold reduction (Fig. 5E). From aligned BAM files to chromosome-arm calls, the average wall-clock processing time per-sample was 0.5 min for FFPE-CUTAC and 46 min for WGS, representing a 92-fold reduction (Fig. 5F). Thus, at 1-Mb resolution, FFPE-CUTAC reproduced WGS-defined chromosome-arm states with 96.0% concordance while substantially reducing long-term data storage and downstream processing time at scale.

## Discussion

This cross-assay benchmark studies FFPE-CUTAC as a robust DNA-based chromatin assay that captures RNAPII-associated regulatory activity and chromosome-arm copy-number information from FFPE tissue (Table 2). FFPE RNA-seq showed collection-year-associated declines in read composition, transcript coverage uniformity, insert length, and gene detection. It also showed lower TIN and less uniform gene-body coverage than matched FF RNA-seq, although these differences also reflect distinct library-enrichment strategies. In contrast, FFPE-CUTAC showed no systematic specimen-age-associated decline in genomic-feature composition, fragment length, gene-associated signal, or peak-level metrics. At matched sequencing depth, FFPE-CUTAC covered a broader genomic space that extended from annotated genes to non-coding regulatory elements. FFPE-CUTAC-favored genes included replication-coupled histone genes, lncRNA loci, and transcription-factor genes that were weakly represented by RNA-seq. FFPE-CUTAC also detected RNAPII-associated activity at a greater fraction of high-confidence transcribed enhancer loci. These genes had shorter reference mRNA half-lives, consistent with FFPE-CUTAC being less constrained by mature RNA stability and recovery because it measures RNAPII occupancy rather than steady-state RNA abundance.

**Table 2:** Genomic features recoverable from FFPE samples across profiling assays.

| Genomic Features from FFPEs | CUTAC | RNA-seq | WGS |
| --- | --- | --- | --- |
| Coding SNPs |  | ✓ | ✓ |
| Fusion proteins |  | ✓ | ✓ |
| Copy number variation | ✓ | ✓ | ✓ |
| Gene expression | ✓ | ✓ |  |
| Regulatory mutations | ✓ |  | ✓ |
| Promoter/Enhancer activity | ✓ |  |  |
| microRNA transcription | ✓ |  |  |
| Hypertranscription | ✓ |  |  |
| Histone gene transcription | ✓ |  |  |

The shared gene-level signal supported integration of FFPE-CUTAC with existing RNA-seq reference cohorts. In meningioma, FFPE-CUTAC aligned closely with FF RNA-seq and retained patient-level correspondence and disease-relevant structure defined by WHO grade, chromosome 22q status, the NF2/Merlin signature, and DNA methylation groups. When FF RNA-seq was unavailable, FFPE RNA-seq also supported integration, although matched-pair alignment was weaker and more variable. In breast cancer, matched FFPE-CUTAC and FFPE RNA-seq profiles aligned within a shared embedding that retained ER^+^/PR*^−^*/HER2*^−^*, HER2^+^, and TNBC subtype structure. These findings support the feasibility of projecting archival FFPE-CUTAC profiles into existing RNA-seq disease maps for molecular stratification.

Because FFPE-CUTAC sequences DNA fragments, each library also retained genome-wide copy number dosage information. At 1-Mb resolution, FFPE-CUTAC achieved 96.0% overall concordance with WGS-defined chromosome-arm states, exceeding CaSpER-based inference from either RNA-seq assay. Concordance between FFPE-CUTAC and WGS was consistently high across patients and chromosome arms. Relative to WGS, FFPE-CUTAC reduced the average data storage by approximately 200-fold and chromosome-arm CNA-calling time by 92-fold under the benchmark environment. Thus, a single shallow, low-input FFPE-CUTAC library provides complementary measurements of RNAPII-associated regulatory activity and chromosome-arm aneuploidy.

Together, these capabilities address a central barrier in retrospective cancer genomics. Hospital pathology archives contain extensive FFPE collections linked to diagnosis, treatment, and long-term clinical outcomes. However, nucleic-acid degradation, tissue-input requirements, and assay burden have limited genome-wide profiling at scale. FFPE-CUTAC offers a practical strategy for converting these specimens into integrated regulatory and copy-number profiles. Integration with existing RNA-seq reference maps may extend molecular disease maps to archival cases lacking fresh-frozen tissue and connect their molecular states with clinical annotations and outcomes.

Several limitations remain. Copy-number benchmarking was restricted to chromosome-arm-level aneuploidy in meningioma. At finer genomic scales, shallow FFPE-CUTAC coverage is sparse and susceptible to local technical bias. Improved methods for modeling sparse read depth, correcting local bias, and segmenting heterogeneous tumor profiles may extend detection to large segments and focal events without sacrificing the low-input, shallow-sequencing workflow. Validation in larger cohorts with diverse tumor purity, ploidy, and CNA burden will also be required.

FFPE-CUTAC is an antibody-guided chromatin profiling assay adapted from CUT&Tag and is compatible with minimal tissue input. This property may enable regional microdissection, multiple profiles from a single 5-*µ*m section, and application to tissue microarrays. Regional sampling would retain coarse spatial context and permit comparison of RNAPII-associated regulatory states and chromosome-arm CNAs across tumor regions. With improved copy-number resolution and further validation, these profiles may also support reconstruction of regional clonal evolution and expansion patterns.

More broadly, FFPE-CUTAC provides a framework for retrospective disease mapping. Archival tumors could be profiled and projected into reference spaces linked to histology, genotype, treatment, and outcome. After external and prospective validation, these maps can support molecular stratification, recurrence-risk modeling, and treatment-association studies. The present study establishes technical feasibility potentially leading to clinical utility.

## Conclusions

FFPE-CUTAC provides a single, shallow, low-input assay for recovering complementary RNAPII-associated regulatory and chromosome-arm copy-number information from archival FFPE tissue. It retains broad genomic signal across recent and older specimens, captures regulatory loci underrepresented by RNA-seq, supports integration with existing RNA-seq reference maps, and reproduces WGS-defined chromosome-arm states with 96.0% overall concordance while substantially reducing storage and computational requirements. By combining these measurements in one library, FFPE-CUTAC provides a practical foundation for extending molecular disease maps to clinically annotated FFPE archives, particularly when fresh-frozen tissue and specimen material are limited.

## Methods

### Fresh-frozen RNA sequencing

Total RNA was isolated from fresh-frozen meningioma tissue using the RNeasy Plus Mini Kit (QIAGEN). RNA integrity was assessed using an Agilent 4200 TapeStation and reported as RNA integrity number equivalent (RINe). Samples with RINe values below 5 were excluded from downstream processing. RNA concentration was measured using a DropSense96 spectrophotometer. RNA was normalized to 50 ng/*µ*L, and 500 ng of total RNA was used for library preparation. Stranded poly(A)-selected RNA-seq libraries were prepared using the TruSeq Stranded mRNA Library Prep Kit (Illumina). Libraries were sequenced on an Illumina NovaSeq 6000 using 50-bp paired-end reads. RNA extraction was performed by the Specimen Processing and Research Cell Bank at Fred Hutchinson Cancer Center. Library preparation and sequencing were performed by the Genomics and Bioinformatics Core at Fred Hutchinson Cancer Center.

### FFPE RNA and DNA extraction and sequencing

FFPE specimens were received as 10-*µ*m tissue curls. Two curls were used for each extraction. Genomic DNA and total RNA were isolated from the same tissue sections using the AllPrep DNA/RNA FFPE Kit (QIAGEN). FFPE RNA quality was evaluated using DV200. A DV200 value above 30% was targeted for library preparation. Libraries that failed post-preparation quality control were excluded from downstream analysis. Stranded RNA-exome libraries were prepared from FFPE total RNA using the TruSeq RNA Exome workflow (Illumina). This workflow uses random-primed cDNA synthesis followed by sequence-specific hybrid capture of predefined coding transcript regions. Whole-genome sequencing libraries were prepared from a target input of 100 ng of FFPE-derived genomic DNA using the xGen cfDNA and FFPE DNA Library Prep Kit (Integrated DNA Technologies). Libraries were sequenced on an Illumina and sequencing targeted a mean genome-wide depth of 120*×*.

### FFPE-CUTAC library preparation and sequencing

FFPE-CUTAC was performed according to the protocol deposited at protocols.io (https://www.protocols.io/view/cutac-for-ffpes-14egn292zg5d/)^3^. Briefly, one FFPE tissue section was used for each experiment without prior bulk nucleic acid extraction. Sections were deparaffinized and subjected to antigen retrieval. RNA polymerase II was targeted using an antibody against the Ser5-phosphorylated C-terminal domain. Protein A–Tn5 was tethered to antibody-bound RNAPII, and tagmentation preferentially released DNA fragments of approximately 120 bp adjacent to RNAPII-occupied loci. Libraries were amplified for 13 PCR cycles. Libraries with insufficient amplified DNA yield, as assessed using an Agilent TapeStation, were excluded from downstream sequencing and analysis.

### Tissue cohorts, read alignment, and genomic annotations

Two FFPE tumor cohorts were analyzed. The meningioma cohort comprised tissue from 30 patients collected between the year of 2017 and 2023. It included 36 FFPE-CUTAC samples, 30 FFPE RNA-seq samples, 17 FF RNA-seq samples, and 30 FFPE WGS samples. The breast cancer cohort comprised tissue from 15 patients collected between 1999 and 2012. It included 25 FFPE-CUTAC samples from 13 patients and 15 FFPE RNA-seq samples. All sequencing reads were aligned to the hg19 reference genome. FFPE-CUTAC reads were aligned with Bowtie 2 (v2.4.4) using `--very-sensitive-local`, `--soft-clipped-unmapped-tlen`, and `--dovetail`. FFPE and FF RNA-seq reads were aligned with STAR (v2.7.11b) using `--twopassMode Basic` and `--quantMode TranscriptomeSAM GeneCounts`. FFPE WGS reads were aligned with BWA-MEM in BWA (v0.7.17) using default parameters. Alignment files and genomic intervals were processed using SAMtools (v1.19.2), BEDTools (v2.30.0) for peak and intermediate interval processing, BEDTools (v2.31.0) for genomic binning, and Sambamba (v1.0.1) for window-level read-depth summarization. Gene-level analyses used GENCODE v19, corresponding to Ensembl release 74 on GRCh37. RefSeq transcript models were obtained from the UCSC hg19 `refGene` table. Candidate cis-regulatory element (cCRE) annotations were obtained from the ENCODE SCREEN hg19 `encodeCcreCombined` track.

### FFPE-CUTAC peak calling, FRiP, and Shannon entropy

FFPE-CUTAC peaks were called with SEACR (v1.3)^118^ in stringent mode using a numeric threshold of 0.10, which retained the top 10% of candidate regions ranked by total signal. Sequencing depth was summarised as the number of aligned reads or fragments per library, depending on the assay-specific counting unit. Genome-coverage breadth was calculated after downsampling each library to 5 million aligned reads or fragments and was defined as the percentage of analysed hg19 bases covered by at least one read or fragment. Peak-calling saturation was assessed by downsampling each FFPE-CUTAC library to 0.5, 1, 2, 4, 6, 8, and 10 million aligned fragments, together with the full-depth data. Peak calling was repeated at each depth using the same SEACR parameters. The fraction of fragments in peaks (FRiP) was defined as the percentage of aligned fragments overlapping at least one called peak. Peak number and FRiP were recorded at each sequencing depth.

Shannon entropy was calculated per library from raw feature-count distributions. RNA-seq entropy was calculated across genes. FFPE-CUTAC entropy was calculated separately across genes, ENCODE cCREs, and non-overlapping genome-wide 500-bp bins. For feature counts *x_i_*, probabilities were defined as

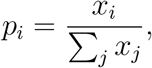

and entropy was calculated as

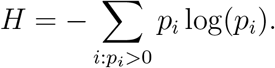

### Genome annotation, RNA integrity, and GC composition

FFPE-CUTAC peaks were annotated with ChIPseeker^119^ (v1.38.0) using `annotatePeak`, `TxDb.Hsapiens.UCSC.hg19.knownGene`, and a transcription start site window of *±*3 kb. RNA-seq read distributions, gene-body coverage, and transcript integrity were evaluated with RSeQC^120^ (v5.0.4) using `read distribution.py`, `geneBody coverage.py`, and `tin.py`, respectively. Transcript integrity number (TIN) per transcript was calculated using a length-stratified RefSeq transcript model comprising approximately 6,000 multi-exon transcripts on chromosomes 1–22 and X. Transcript lengths ranged from 0.5 to 250 kb, with approximately equal numbers sampled from each length decile.

For matched meningioma RNA-seq libraries, the paired TIN difference was calculated for transcripts with non-zero TIN in both FFPE and FF libraries from the same patient:

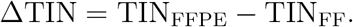

Negative values indicate lower transcript integrity in FFPE than in matched FF RNA-seq. We evaluated associations of transcript-level TIN and ΔTIN with transcript length, exon count, coding-sequence length, spliced-transcript GC content, and mRNA half-life. We also evaluated 5*^′^* and 3*^′^* UTR lengths defined relative to transcript strand. Transcript length, 3*^′^* UTR length, and mRNA half-life were log-transformed. Continuous features were grouped into equal-frequency quintiles. mRNA half-life was additionally grouped into tertiles for age-stratified TIN analyses. TIN was not calculated for FFPE-CUTAC because the metric is defined from RNA-seq coverage across spliced transcript models, whereas FFPE-CUTAC measures RNA polymerase II-associated genomic DNA fragments.

GC content was quantified from approximately 250,000 reads or fragments per library with custom Python scripts using pysam (v0.24.0). For RNA-seq, GC content was calculated from aligned read sequences. For WGS, GC content was calculated from the hg19 reference sequence across the genomic span of each properly paired template. For FFPE-CUTAC, GC content was calculated from the hg19 reference sequence across fragment intervals inferred from Bowtie2 alignments. Observed DNA-fragment GC distributions were compared with those from length-matched random genomic intervals to quantify enrichment relative to genomic background. WGS GC bias was independently evaluated with deepTools (v3.5.5) `computeGCBias` using the hg19 two-bit reference genome. RNA-seq was excluded from the genomic-background comparison because RNA-seq fragments can span splice junctions and therefore do not correspond to a single contiguous genomic interval.

### RNA half-life annotation and transcript-stability analyses

Gene-level RNA half-lives were obtained from RNAdecayCafe v1.1 using `AvgKdegs genes v1.1.csv` from Zenodo record 16884513^100^. RNAdecayCafe reports degradation-rate constants and half-lives estimated from nucleotide-recoding RNA sequencing across eleven human cell lines. Half-life was used in hours. A consensus half-life was computed as the median value across the eleven cell lines for each gene. This procedure yielded half-life estimates for 17,412 genes. Estimates were mapped to GENCODE v19 genes by gene symbol. Because the source measurements were obtained from cultured cell lines rather than the tumour tissues analysed here, they were treated as proxies for relative gene-level RNA stability. Half-life annotations were used in three analyses. First, transcripts with matched gene-level annotations were divided into unstable, intermediate, and stable tertiles for collection-year-associated TIN analyses. Second, log-transformed half-life was included as a standardised covariate in multivariable models of ΔTIN. Third, half-life distributions were compared between FFPE-CUTAC-favored genes and co-detected genes in gene-level comparisons of FFPE-CUTAC with RNA-seq.

### Fragment length and gene-level comparison of FFPE-CUTAC and RNA-seq

Median FFPE-CUTAC fragment length and RNA-seq insert length were calculated for each sample from paired-end alignments. Genes with at least one assigned read or fragment were considered detected. RNA-seq gene detection was based on the unstranded counts in STAR `ReadsPerGene.out.tab`, whereas FFPE-CUTAC gene detection was based on GENCODE (v19) gene-level fragment counts. Associations with tissue collection year were assessed by ordinary least-squares regression and Spearman’s rank correlation.

Within each tumor cohort, RNA-seq reads and FFPE-CUTAC fragments were quantified over GENCODE (v19) genes. Gene-level signals were analyzed as raw counts and log-transformed counts per million (log-CPM), respectively. Genes were classified as FFPE-CUTAC-favored when the raw RNA-seq count was greater than 0 but less than 30 and the raw FFPE-CUTAC count was greater than 30.

### RNA-category annotation and enrichment testing

Co-detected genes were annotated using GENCODE v19 gene biotypes. For the broad classification in Fig. 3D, protein-coding genes were defined by the `protein coding` biotype. Long non-coding RNA genes comprised `lincRNA`, `antisense`, `sense intronic`, `sense overlapping`, `processed transcript`, and `3prime overlapping ncrna`. The small-RNA category comprised `miRNA`, `snRNA`, `snoRNA`, `misc RNA`, and `rRNA`. Immunoglobulin and T-cell receptor gene segments with biotypes beginning with `IG` or `TR` were assigned to the immune V(D)J category before pseudogene classification. Remaining biotypes ending in `pseudogene` were assigned to the pseudogene category.

Transcription factors were defined as protein-coding genes present in either of two reference sets: Human transcription-factor census database (v1.01)^77^ and Gene Ontology term GO:0003700 (i.e., DNA-binding transcription factor activity), including descendant terms from `org.Hs.eg.db`. Genes were matched by upper-case HGNC symbol or version-stripped Ensembl gene ID. Transcription-factor structural-family composition was evaluated among FFPE-CUTAC-favored transcription factors. DNA-binding-domain families were taken from the Human transcription-factor census database^77^ `DBD` class field.

For each category, enrichment in the FFPE-CUTAC-favored gene set was tested using a two-sided Fisher’s exact test. Each 2 *×* 2 contingency table crossed gene category membership (rows) with FFPE-CUTAC-favored or RNA-favored category (columns). Odds ratios and exact 95% confidence intervals were reported. *P* values were adjusted using the Benjamini–Hochberg method separately within each combination of cohort, normalization method, and category scheme.

### cCRE signal, enhancer detection, and stringent eRNA filtering

The cCRE regions was obtained from the ENCODE SCREEN `encodeCcreCombined` track. cCREs were classified as promoter-like signatures, proximal enhancer-like signatures, distal enhancer-like signatures, DNase-H3K4me3 elements, or CTCF-only elements. Intergenic enhancer cCREs were defined as proximal or distal enhancer-like cCREs that did not overlap any GENCODE v19 gene body, yielding 121,335 intervals. For each library, enhancer detection was defined as the percentage of these intervals with at least one assigned read or fragment. FFPE-CUTAC counts were obtained from the cCRE-level fragment-count matrix. For RNA-seq, reads overlapping the same intervals were quantified from STAR-aligned BAM files with `featureCounts` in Subread (v2.0.6). A SAF annotation of the enhancer intervals and paired-end, unstranded counting were used with `-p --countReadPairs -s 0`.

High-confidence eRNA loci were defined as intergenic enhancer cCREs that satisfied three criteria (Supplementary Fig. 4Q). First, potential readthrough transcription was excluded. Expressed genes were defined as genes with a raw RNA-seq count greater than 5 in at least one RNA-seq library. A strand-aware 10-kb region immediately downstream of each expressed gene was then defined. For plus-strand genes, this region extended from gene end +1 bp to gene end +10 kb. For minus-strand genes, it extended from gene start *−*10 kb to gene start *−*1 bp. cCREs overlapping any downstream region were excluded. Second, cCREs were required to overlap the FANTOM5 permissive enhancer atlas on hg19^121^. Third, cCREs were required to show bidirectional RNA transcription. Each candidate cCRE was represented by two SAF features with identical coordinates and opposite strands. Strand-specific RNA-seq fragments were quantified with `featureCounts` using paired-end, reverse-stranded counting with `-p --countReadPairs -s 2`. Counts were summed across RNA-seq libraries separately for each strand. A cCRE was considered bidirectionally transcribed when at least two fragments were assigned to each strand. The intersection of these three filters yielded 220 high-confidence eRNA loci. For each library, eRNA-locus detection was defined as the percentage of these loci with at least one assigned read or fragment.

### Shared embedding of FFPE-CUTAC and RNA-seq by CCA integration

FFPE-CUTAC and RNA-seq libraries were integrated into a shared sample-level space using the canonical correlation analysis (CCA) anchor-based integration workflow in Seurat (v5.0.0). Each observation represented one bulk sequencing library. RNA-seq input comprised STAR gene-level counts. When multiple RNA-seq datasets were combined, raw count matrices were adjusted for study- or cohort-associated batch effects using ComBat-seq, implemented as `ComBat seq` in `sva` (v3.50.0).

FFPE-CUTAC fragments were counted over a LINE-extended GENCODE v19 gene intervals. Each interval was extended from the annotated 3*^′^* end in the direction of transcription to the nearest downstream LINE element or until an intervening gene boundary was reached. These intervals were designed to capture downstream read-through RNAPII signal. Genes failing the RNA-seq count filter were removed, and both matrices were restricted to genes shared between modalities. Within each modality, counts were normalized with `LogNormalize` using `scale.factor = 1e6`, yielding log(1 + CPM) values. Variable features were selected with `FindVariableFeatures` using `selection.method = “vst”`. normalized values were centred and scaled gene-wise across libraries with `ScaleData`. An initial 35-component PCA was computed for each modality with `RunPCA(npcs = 35)`. Integration features were selected with `SelectIntegrationFeatures`. Cross-modality anchors were identified with `FindIntegrationAnchors(reduction = “cca”)` using the dataset-specific `k.anchor` values reported below. Candidate anchors were mutual nearest-neighbour library pairs in the CCA space and were filtered and scored by Seurat. Integrated values were generated with `IntegrateData` using the corresponding `k.weight` values. The integrated assay was centred and scaled, reduced by PCA, and visualised with UMAP using PCs 1 through `pca num`, `n.neighbors = 15`, and `min.dist = 1e-4`.

Same-patient correspondence was evaluated from the stored anchor pairs. For each FFPE-CUTAC library, its anchored RNA-seq partners were identified. A pairing was classified as correct when both libraries originated from the same patient. Matched-pair distance was calculated as the Euclidean distance between FFPE-CUTAC and RNA-seq libraries in the first 50 PCs of the corresponding integrated assay. Distances were compared only within the same integrated object. UMAP coordinates were used only for visualisation.

### FFPE-CUTAC copy-number alteration calling at chromosome arm level

Arm-level CNAs were inferred from broad genomic variation in FFPE-CUTAC read depth (Fig. 5A). FFPE-CUTAC alignments were coordinate-sorted and indexed with samtools. Mean read depth was summarized in non-overlapping 1-Mb hg19 windows with sambamba. The 1-Mb resolution was used for the primary analysis to stabilize depth estimates at the shallow sequencing depths of FFPE-CUTAC. Bin-level depth profiles were processed with QDNAseq (v1.38.0) using hg19 bin annotations. Unreliable bins were removed with `applyFilters(residual = TRUE, blacklist = TRUE)`. Joint GCcontent and mappability bias was estimated by two-dimensional loess regression with `estimateCorrection` and removed with `correctBins`. `normalizeBins` scaled each profile to its genome-wide median, yielding within-sample relative depth ratios. `smoothOutlierBins` reduced isolated outlier bins. The normalized ratios were segmented by circular binary segmentation using QDNAseq `segmentBins(transformFun = “none”)` and DNAcopy (v1.76.0). Segment-level ratios were recentered with `normalizeSegmentedBins`.

Chromosome-arm intervals were defined from centromeric gap coordinates in the UCSC hg19 gap table, with centromeric regions excluded. Corrected bin-level log_2_ ratios were segmented by circular binary segmentation (CBS) using QDNAseq `segmentBins(transformFun = “none”)` and DNAcopy (v1.76.0). Segmented bins were normalized with `normalizeSegmentedBins`. A segment was classified as gained when its normalized depth ratio exceeded 2^0.5^ and lost when its ratio was below 2*^−0.5^*. These thresholds correspond to log_2_ ratios greater than 0.5 and less than *−*0.5, respectively. An arm was called gain when gained segments covered at least 50% of the arm interval and loss when lost segments met the same criterion. All remaining arms were classified as intact. The five acrocentric short arms, 13p, 14p, 15p, 21p, and 22p, were excluded, leaving 39 evaluable autosomal chromosome arms. The resulting patient-by-arm categorical matrix was benchmarked directly against the WGS arm-level calls.

Resolution sensitivity was evaluated using 100-kb, 500-kb, 1-Mb, 2-Mb, and 5-Mb profiles. The 2-Mb and 5-Mb profiles were generated by aggregating corrected 1-Mb bins. Technical reproducibility was assessed as arm-call concordance across replicate FFPE-CUTAC libraries from the same patient.

### WGS depth-based copy-number alteration calling

WGS copy-number profiles were used as the gold standard for arm-level benchmarking analysis. Coordinate-sorted and indexed WGS BAM files were filtered with samtools using `-q 20 -F 0xF00`. This retained reads with mapping quality at least 20 and removed secondary alignments, supplementary alignments, duplicate reads, and QC-failed reads. Read depth was summed in fixed 100-kb genomic windows with bedtools. Poorly mappable or blacklisted bins were removed with QDNAseq (v1.38.0) using `applyFilters(residual = TRUE, blacklist = TRUE)`.

An allele-specific WGS pipeline, ASCAT (v3.2.0), was applied to WGS BAM files using 1000 Genomes hg19 SNP loci and alleles. Allele counts were generated with *canceritallelecount* (v4.3.0) through the ASCAT high-throughput sequencing workflow. ASCAT corrected logR values for GC content and replication timing, inferred germline heterozygous sites, co-segmented logR and B-allele frequency with allele-specific piecewise constant fitting, and estimated tumor purity and ploidy. Allele-specific copy-number segments were reduced to arm-level gain or loss calls if more than 50% of the arm length is deemed gain or loss. We have also adapted the same processing procedure designed for FFPE-CUTAC to WGS to generate the log_2_ signal-to-baseline ratio profiles for visual comparison.

### RNA-seq expression-based copy-number alteration calling

FFPE RNA-seq and FF RNA-seq CNAs were inferred with CaSpER (v0.2.0). The two RNA-seq assays used the same expression-based pipeline and differed only in their expression input. Raw gene counts were supplied to CaSpER with `matrix.type = “raw”`, `log.transformed = FALSE`, and `expr.cutoff = 10`. B-allele frequency was extracted from RNA-seq BAM files using BAFExtract. BAFExtract has no released version tag and was treated as the CaSpER companion source build. Pileups were generated against hg19 with `generate compressed pileup per SAM`. Single-nucleotide variants were retained with minimum coverage of 20, minimum alternate count of 4, and minimum minor allele frequency of 0.1. BAF files were imported into CaSpER with `readBAFExtractOutput(sequencing.type = “bulk”)`. CaSpER objects were constructed with `CreateCasperObject`. Parameters were `cnv.scale = 3`, `loh.scale = 3`, `window.length = 50`, `length.iterations = 50`, `filter = “median”`, `matrix.type = “raw”`, `expr.cutoff = 10`, `sequencing.type = “bulk”`, `genomeVersion = “hg19”`, and `log.transformed = FALSE`. CaSpER internally normalized expression to the external reference set and recursively median-smoothed the expression signal across three genomic scales. BAF was processed across three LOH scales. Segmentation was performed with `runCaSpER(method = “iterative”, removeCentromere = TRUE)`. Large-scale events were extracted with `extractLargeScaleEvents`. This produced sample-by-arm matrices of gain, loss, and neutral status. Two thresholds were analyzed independently. The primary default threshold was 0.75. A more stringent threshold of 0.9 was also reported. The same five acrocentric short arms were removed, leaving the same 39 chromosome arms used for WGS and FFPE-CUTAC benchmarking.

### Statistical analysis and data visualisation

Statistical analyses were performed in R (v4.3.2). Ordinary least-squares regression and Spearman correlation were used to assess trends with tissue collection year. Two-sided Wilcoxon rank-sum tests were used for group comparisons unless otherwise specified. Fisher’s exact tests were used for category enrichment. P values were adjusted with the Benjamini-Hochberg method within the relevant analysis stratum. Drivers of FFPE-specific RNA integrity loss were modeled by multiple linear regression with standardised coefficients. The response and predictors were scaled to zero mean and unit variance. Predictors included log transcript length, log 3*^′^*UTR length, GC content, and log half-life. Because transcript length and 3*^′^*UTR length are correlated, multivariable length coefficients were interpreted together with univariable estimates. Plots were generated with ggplot2 (v4.0.0) and assembled with ggpubr (v0.6.1).

### Computational environment and benchmark measurements

Data storage and processing-time benchmarks were performed on a single shared compute node equipped with two AMD EPYC 9554 64-core processors, providing 128 physical CPU cores and 256 logical threads. The node had 3.0 TiB of RAM and ran Red Hat Enterprise Linux 9.7 with Linux kernel 5.14.0. Aligned sequencing files were accessed from a shared network filesystem. Because all samples for a given benchmark step could be processed concurrently on this node, runtime was reported as per-sample wall-clock time under parallel whole-cohort processing. This measurement reflects practical cohort-scale throughput on the shared server environment, rather than isolated single-sample runtime.

## Data availability

FFPE-CUTAC sequencing data analysed in this study are available from the Gene Expression Omnibus (GEO) under accession GSE261351. Associated processed data are available from Zenodo at https://doi.org/10.5281/zenodo.13138686. Meningioma fresh-frozen RNA-seq data were obtained from the GEO under accession GSE252291^58^. FFPE RNA-seq data analysed in this study are available from Zenodo at `ZENODO_FFPE_RNASEQ_DOI`. Whole-genome sequencing data used for copy-number benchmarking are available from Zenodo at `ZENODO_WGS_DOI`.

## Code availability

Custom code used for data processing, modelling, statistical analysis, and figure generation will be made available at https://github.com/CompBioWizard/FFPECUTAC-RNA-WGS. The repository includes the analysis scripts and documentation required to reproduce the computational analyses and figures presented in this study.

## Acknowledgments

This work was supported by the National Institutes of Health NIH/NHGRI grant (R00HG012797) and CCSG New Faculty Award supported by the NIH/NCI grant (P30CA016672) to Y.Z.

## Author Contributions

Y.Z., S.H., and K.A. conceived the project. Y.Z. designed the study. Y.N. and Q.X. performed the benchmarking analyses and developed the FFPE-CUTAC-specific method for copy-number alteration detection. C.Y.M. and Y.X. contributed to assay design, FFPE-CUTAC profiling and experimental benchmarking. Y.N. and A.P. conducted the literature review of comparative profiling and copy-number alteration detection methods. A.Z. and E.H. provided the meningioma and breast cancer RNA-seq and whole-genome sequencing datasets. N.K. and R.P. provided guidance on RNA-seq analysis and clinical interpretation.

---

**Supplementary Figure 1.**
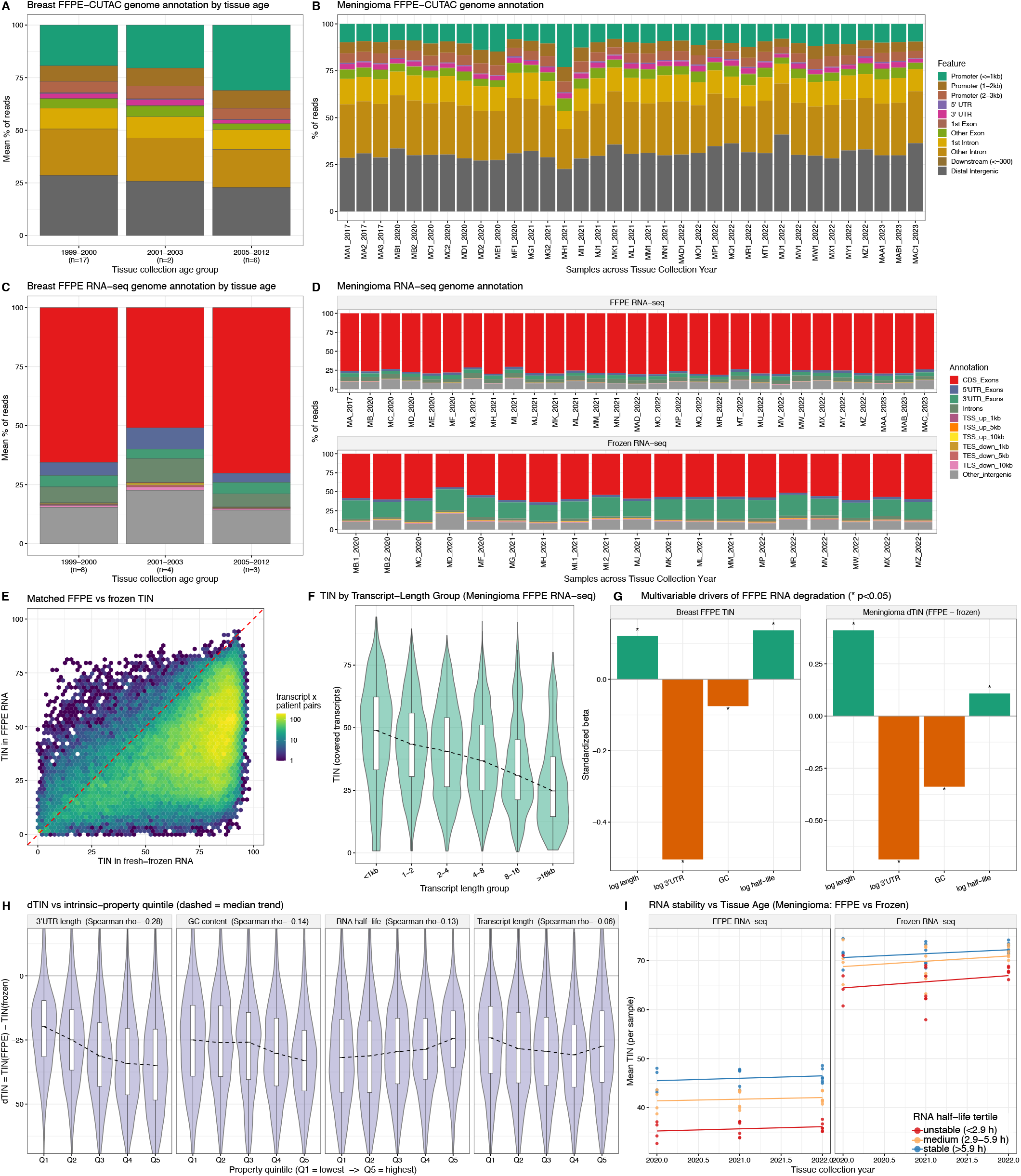
Genome annotation and transcript integrity across FFPE and fresh-frozen specimens. **A.** Mean distribution of breast cancer FFPE-CUTAC reads across genomic-feature categories, stratified by tissue collection period. Stacked bars show category percentages, and the number of libraries is indicated for each period. **B.** Per-sample distribution of meningioma FFPE-CUTAC reads across genomic-feature categories. Samples are ordered by tissue collection year. **C.** Mean distribution of breast cancer FFPE RNA-seq reads across genomic-feature categories, stratified by tissue collection period. The number of libraries is indicated for each period. **D.** Per-library distribution of meningioma RNA-seq reads across genomic-feature categories, shown separately for FFPE and FF RNA-seq. Samples are ordered by tissue collection year. FF replicate libraries are distinguished by the suffixes .1 and .2. **E.** Matched transcript integrity number (TIN) values from meningioma FFPE and FF RNA-seq. Each hexagonal bin summarizes transcript-by-patient pairs with TIN measured in both assays. Bin color indicates the number of pairs on a logarithmic scale, and the red dashed line marks the identity line. dTIN was defined as TIN_FFPE_*−*TIN_FF_. A total of 79,669 transcript-by-patient pairs were analyzed, with a median dTIN of *−*28.1. **F.** TIN distributions among transcripts in meningioma FFPE RNA-seq, stratified by transcript-length group. The dashed line connects group medians. **G.** Standardized coefficients from multivariable linear models of RNA-integrity metrics. Models used breast cancer FFPE RNA-seq TIN or meningioma dTIN as the response. Predictors included log-transformed transcript length, log-transformed 3*^′^* UTR length, GC content, and log-transformed RNA half-life. Responses and predictors were standardized before model fitting. Asterisks indicate coefficients with *P <* 0.05. **H.** Meningioma dTIN distributions stratified by quintiles of 3*^′^* UTR length, GC content, RNA half-life, and transcript length. Q1 and Q5 denote the lowest and highest quintiles, respectively. Facet titles report Spearman correlation coefficients between each property and dTIN. Dashed lines connect median dTIN values across quintiles. **I.** Mean TIN per sample by RNA half-life category for meningioma FFPE and FF RNA-seq as a function of tissue collection year. RNA half-life categories were defined using RNAdecayCafe^100^ as unstable (*<* 2.9 h), medium (2.9–5.9 h), or stable (*>* 5.9 h). Lines show ordinary least-squares fits within each category. In violin plots, widths represent kernel-density estimates. Internal box plots show medians and first and third quartiles, with whiskers extending to the most extreme values within 1.5 times the interquartile range.

**Supplementary Figure 2.**
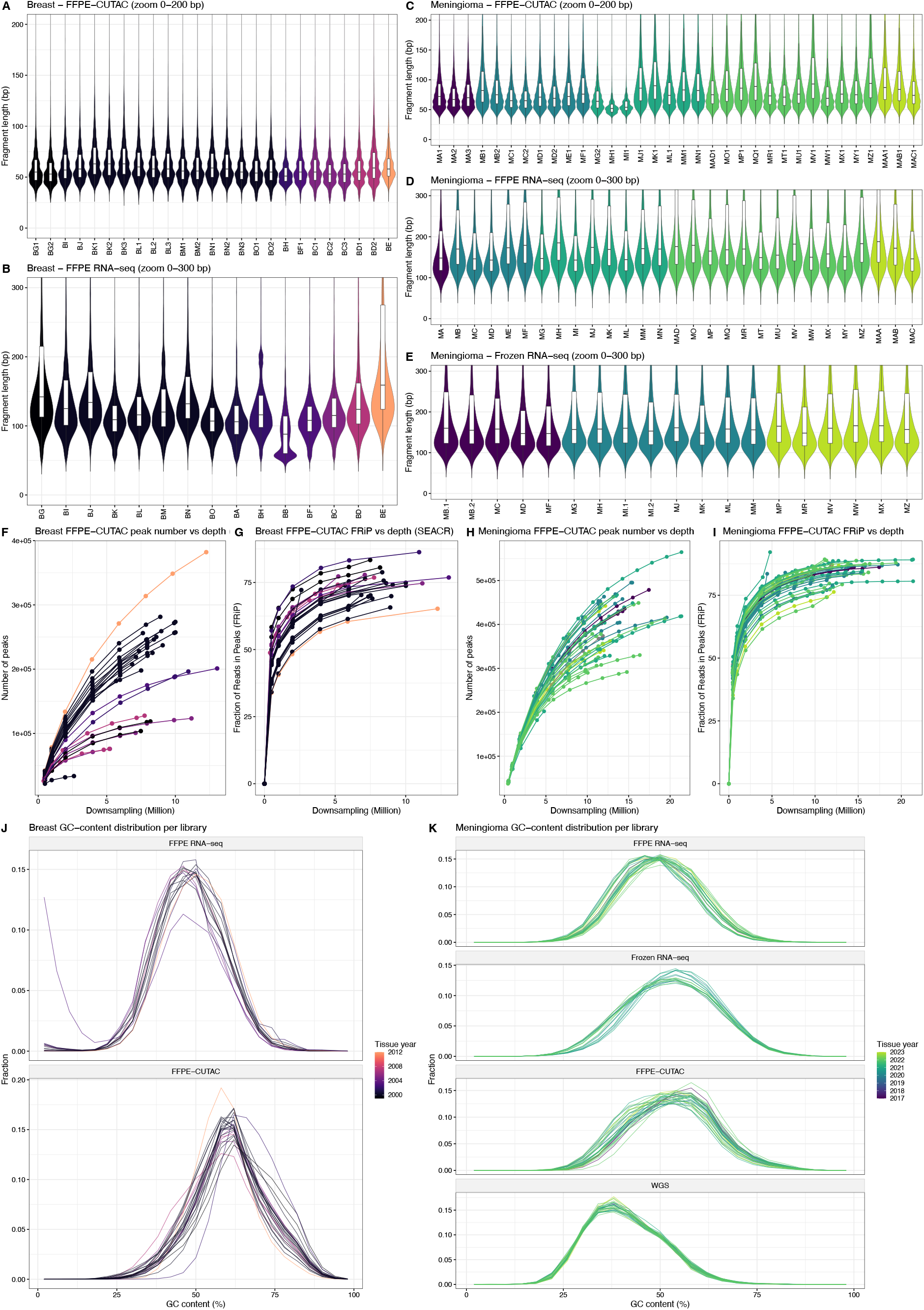
Fragment and insert lengths, FFPE-CUTAC peakcalling saturation, and GC-content distributions. **A.** Fragment-length distributions for breast cancer FFPE-CUTAC libraries. **B.** Insert-length distributions for breast cancer FFPE RNA-seq libraries. **C.** Fragment-length distributions for meningioma FFPE-CUTAC libraries. **D.** Insert-length distributions for meningioma FFPE RNA-seq libraries. **E.** Insert-length distributions for meningioma FF RNA-seq libraries. In **A–E**, libraries are ordered by tissue collection year. Violin widths represent kernel-density estimates and contain overlaid box plots. The displayed y-axis ranges are restricted to 0–200 bp for FFPE-CUTAC and 0–300 bp for RNA-seq. **F.** Peak-calling saturation for breast cancer FFPE-CUTAC libraries, quantified as the number of peaks called at the SEACR top 10% threshold across downsampled sequencing depths. **G.** Fraction of reads in peaks (FRiP) for breast cancer FFPE-CUTAC libraries across the same downsampled depths. **H.** Peak-calling saturation for meningioma FFPE-CUTAC libraries, quantified as the number of peaks called at the SEACR top 10% threshold across downsampled sequencing depths. **I.** FRiP for meningioma FFPE-CUTAC libraries across the same downsampled depths. **J.** GC-content distributions for breast cancer FFPE RNA-seq and FFPE-CUTAC libraries. **K.** GC-content distributions for meningioma FFPE RNA-seq, FF RNA-seq, FFPE-CUTAC, and FFPE whole-genome sequencing libraries. In **J** and **K**, each line represents one library.

**Supplementary Figure 3.**
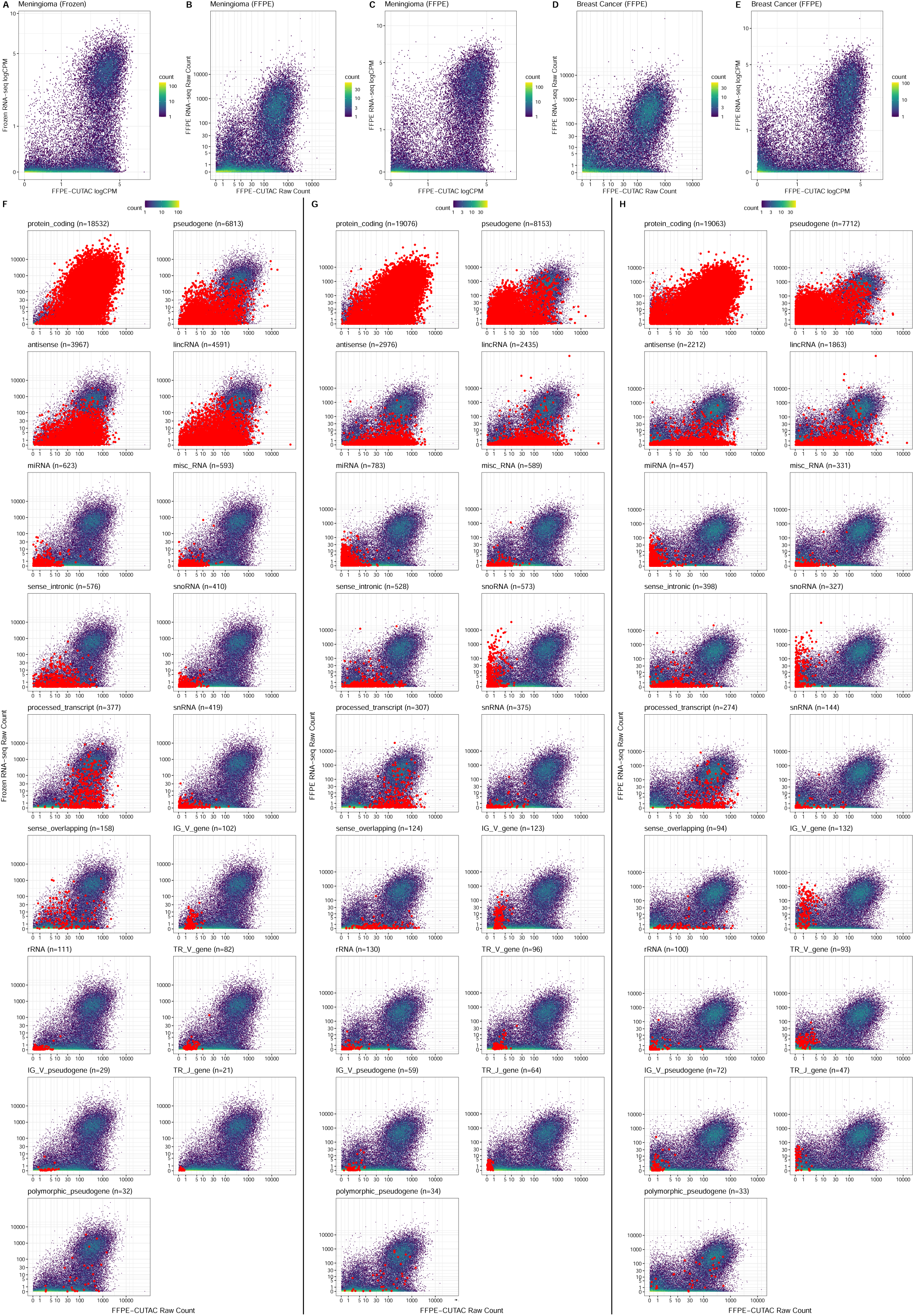
Gene-level concordance between FFPE-CUTAC and RNA-seq across cohorts, normalizations, and gene biotypes. **A.** Mean per-gene logCPM values for meningioma FFPE-CUTAC and FF RNA-seq across matched libraries. **B.** Mean per-gene raw counts for meningioma FFPE-CUTAC and FFPE RNA-seq across matched libraries. **C.** Mean per-gene logCPM values for meningioma FFPE-CUTAC and FFPE RNA-seq across matched libraries. **D.** Mean per-gene raw counts for breast cancer FFPE-CUTAC and FFPE RNA-seq across matched libraries. **E.** Mean per-gene logCPM values for breast cancer FFPE-CUTAC and FFPE RNA-seq across matched libraries. **F.** Gene-biotype-specific raw-count comparison grid for meningioma FFPE-CUTAC versus FF RNA-seq. Each subpanel shows the full gene-level density background, with genes from the indicated GENCODE biotype highlighted in red. Subpanel titles indicate the biotype and the number of genes assigned to that biotype. **G.** Gene-biotype-specific raw-count comparison grid for meningioma FFPE-CUTAC versus FFPE RNA-seq, displayed as in **F**. **H.** Gene-biotype-specific raw-count comparison grid for breast cancer FFPE-CUTAC versus FFPE RNA-seq, displayed as in **F**. In density plots, color indicates the number of genes per bin. Raw-count panels use pseudolog axes with breaks at 0, 1, 5, 10, 30, 100, 1,000, and 10,000. logCPM panels use linear axes with breaks at 0, 1, 5, and 10.

**Supplementary Figure 4.**
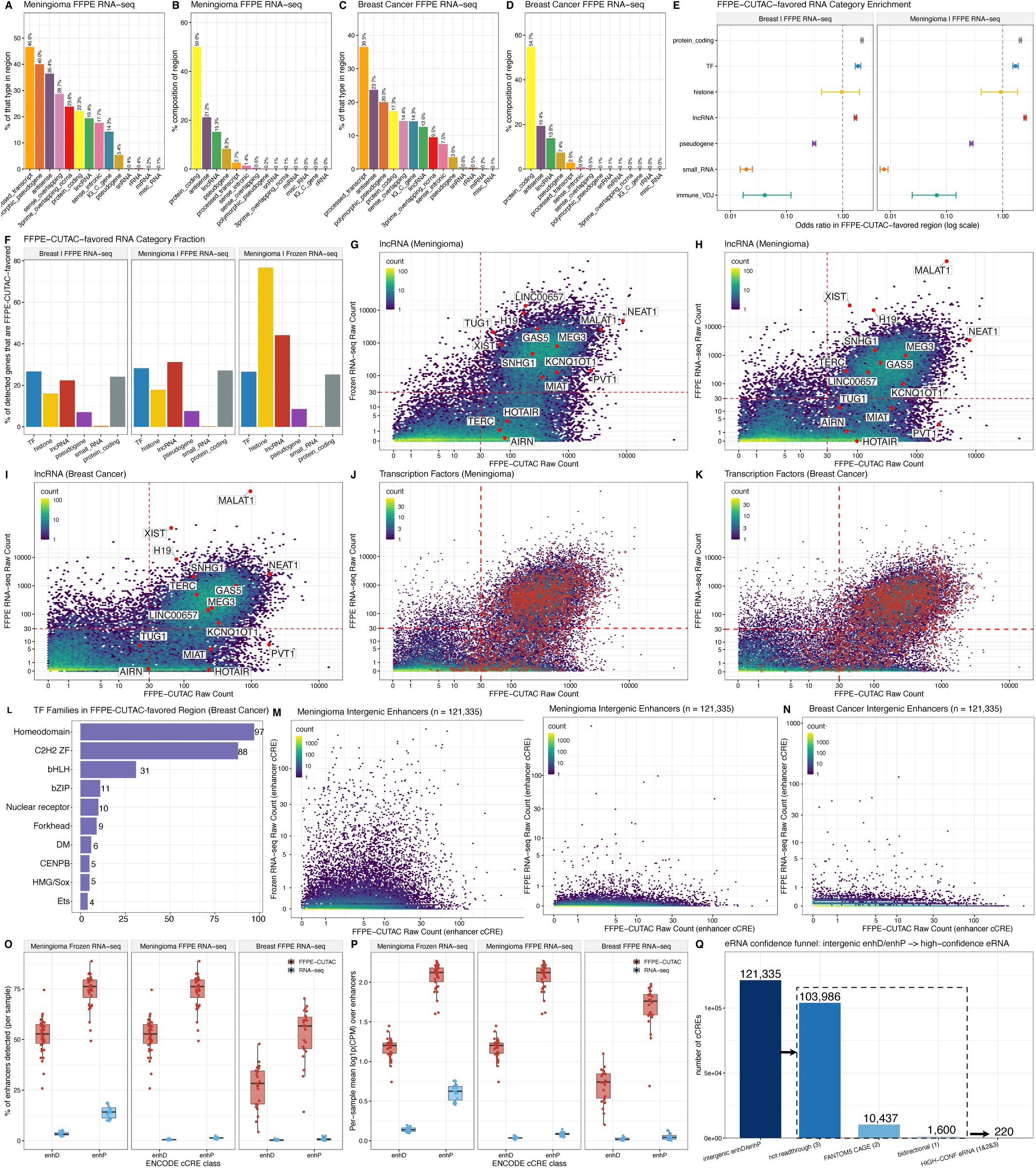
Gene categories and enhancer features enriched among FFPE-CUTAC-favored signals. **A.** Percentage of genes within each GEN-CODE biotype classified as FFPE-CUTAC-favored in the meningioma FFPE RNA-seq comparison. **B.** GENCODE biotype composition of the FFPE-CUTAC-favored gene set in the meningioma FFPE RNA-seq comparison. **C.** Percentage of genes within each GENCODE biotype classified as FFPE-CUTAC-favored in the breast cancer FFPE RNA-seq comparison. **D.** GENCODE biotype composition of the FFPE-CUTAC-favored gene set in the breast cancer FFPE RNA-seq comparison. **E.** Enrichment of broad RNA categories in the FFPE-CUTAC-favored gene set for breast cancer and meningioma FFPE RNA-seq comparisons. Points indicate odds ratios from Fisher’s exact tests relative to the shared gene set, horizontal bars indicate 95% confidence intervals, and vertical dashed lines indicate an odds ratio of 1. **F.** Percentage of detected genes in each RNA category classified as FFPE-CUTAC-favored across the breast cancer FFPE RNA-seq, meningioma FFPE RNA-seq, and meningioma FF RNA-seq comparisons. **G.** Selected nuclear-retained lncRNAs highlighted and labeled on the meningioma FFPE-CUTAC versus FF RNA-seq raw-count density plot. **H.** Selected nuclear-retained lncRNAs highlighted and labeled on the meningioma FFPE-CUTAC versus FFPE RNA-seq raw-count density plot. **I.** Selected nuclear-retained lncRNAs highlighted and labeled on the breast cancer FFPE-CUTAC versus FFPE RNA-seq raw-count density plot. **J.** Transcription factor genes highlighted on the meningioma FFPE-CUTAC versus FFPE RNA-seq raw-count density plot. **K.** Transcription factor genes highlighted on the breast cancer FFPE-CUTAC versus FFPE RNA-seq raw-count density plot. **L.** Structural-family composition of transcription factors in the breast cancer FFPE-CUTAC-favored gene set. Bar-end labels indicate transcription factor counts in FFPE-CUTAC-favored gene set. **M.** Raw-count density plots for intergenic enhancer-like cCREs in meningioma, comparing FFPE-CUTAC with FF RNA-seq (left) and FFPE RNA-seq (right). **N.** Raw-count density plot for intergenic enhancer-like cCREs in breast cancer, comparing FFPE-CUTAC with FFPE RNA-seq. **O.** Per-sample percentage of detected intergenic enhancer-like cCREs, shown by assay, cCRE class, and cohort comparison. **P.** Per-sample mean log_1_*_p_*(CPM) across intergenic enhancer-like cCREs, shown by assay, cCRE class, and cohort comparison. **Q.** Filtering scheme used to define high-confidence enhancer RNAs from intergenic enhD and enhP cCREs. Bars show the number of cCREs retained after each step, including all intergenic enhD/enhP cCREs, removal of readthrough candidates, overlap with FANTOM5 CAGE-supported enhancers, bidirectionality filtering, and the final high-confidence enhancer RNA set. In density plots, color indicates the number of features per bin. In box plots, center lines denote medians, box limits denote the first and third quartiles, and whiskers extend to the most extreme values within 1.5 times the interquartile range. Points represent individual libraries.

**Supplementary Figure 5.**
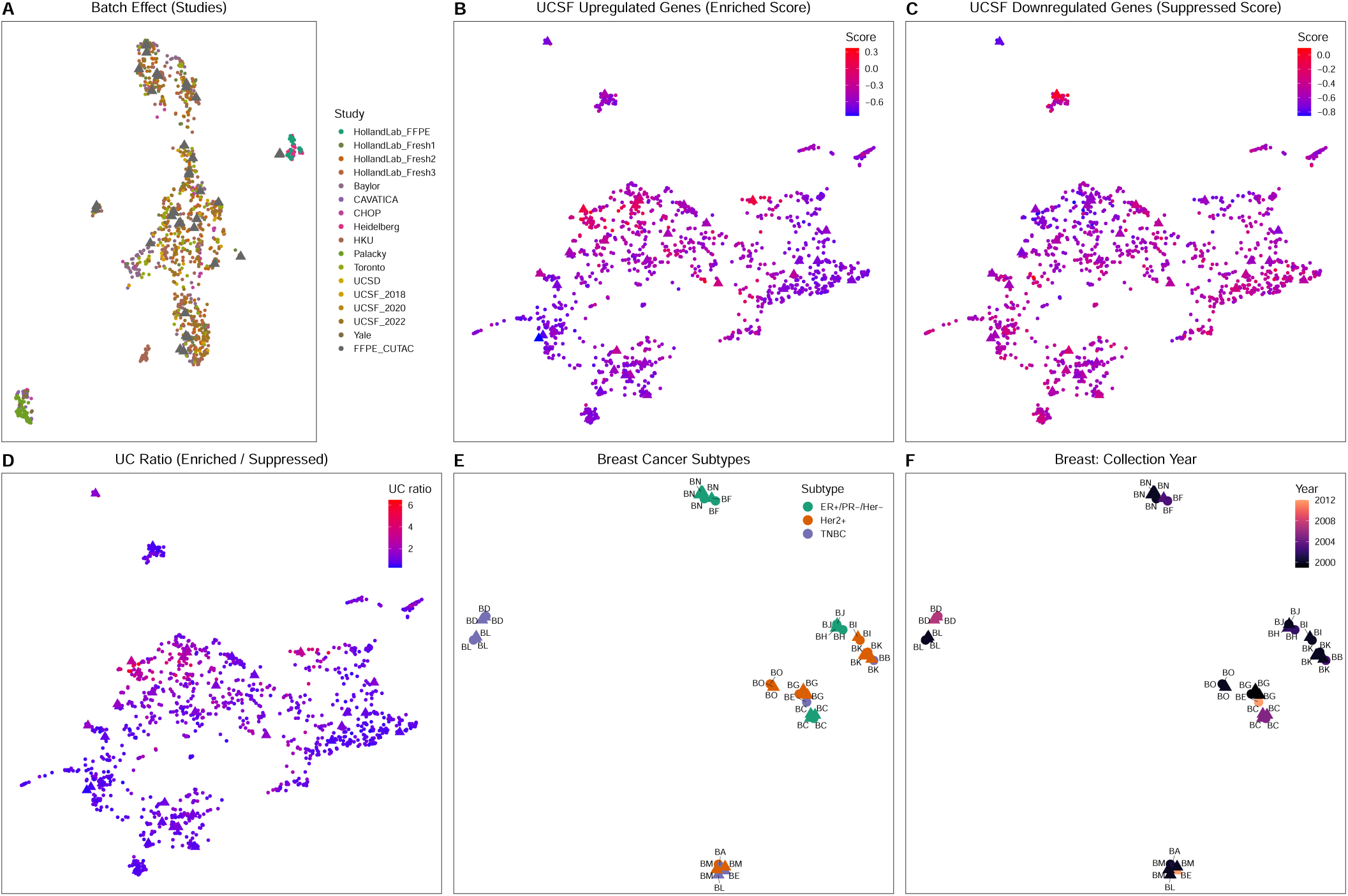
Integrated disease-map embeddings of FFPE-CUTAC and RNA-seq libraries. **A.** Integrated meningioma embedding of FFPE-CUTAC, FF RNA-seq, and FFPE RNA-seq libraries, colored by study of origin. **B.** Same meningioma embedding, colored by the UCSF upregulated-gene enriched score calculated from the 34 meningioma-associated genes^108^. **C.** Same meningioma embedding, colored by the UCSF downregulated-gene suppressed score^108^. **D.** Same meningioma embedding, colored by the UC ratio, defined as (UCSF enriched score +1)/(UCSF suppressed score +1)^108^. **E.** Integrated breast cancer embedding of FFPE-CUTAC and FFPE RNA-seq libraries, colored by breast cancer subtype. **F.** Same breast cancer embedding, colored by tissue collection year.

**Supplementary Figure 6.**
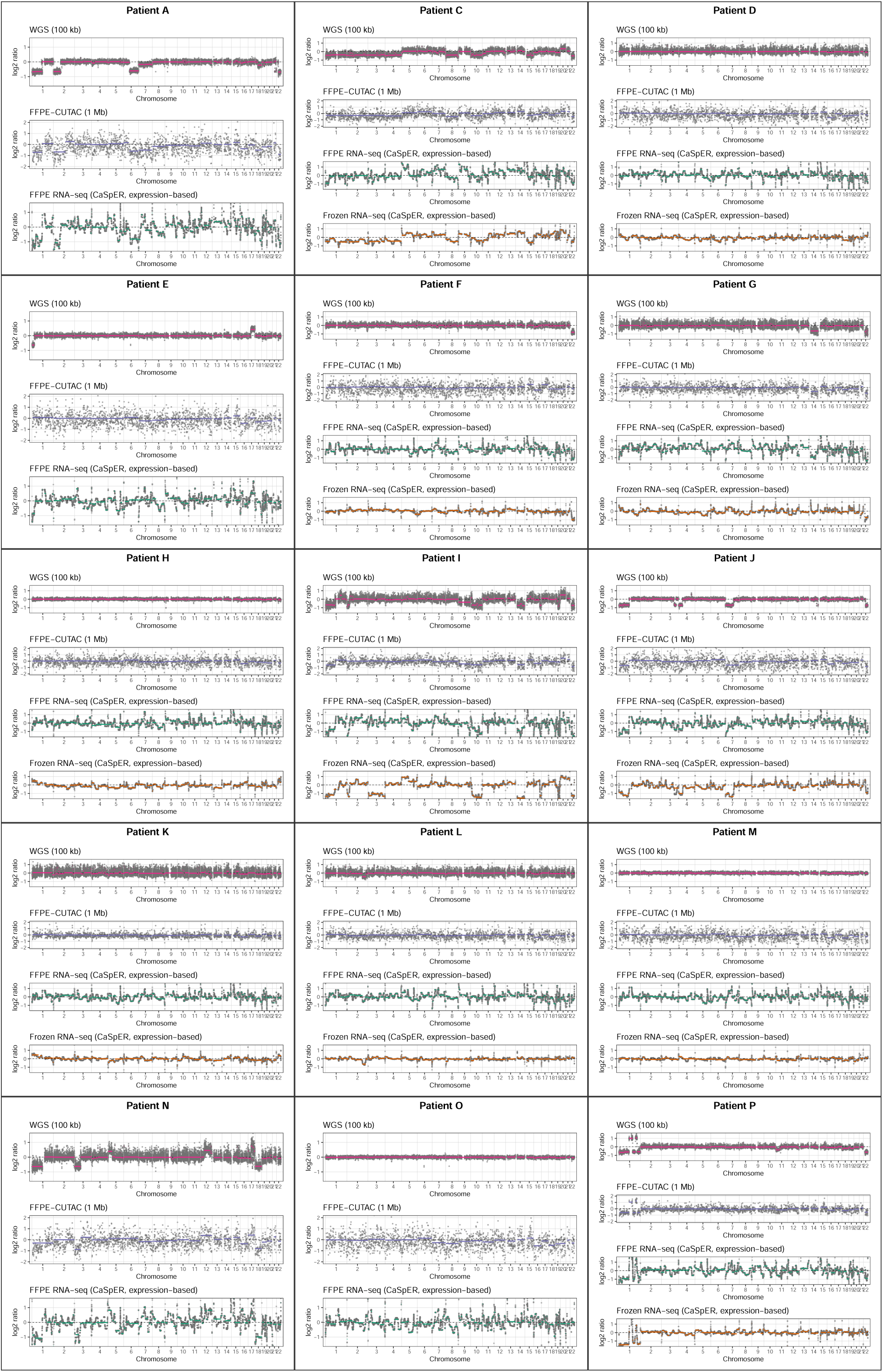
Per-patient copy-number profiles comparison across FFPE WGS, FFPE-CUTAC, and RNA-seq. Copy-number profiles are shown for 15 meningioma patients: A, C, D, E, F, G, H, I, J, K, L, M, N, O, and P. Patient B is presented as the representative example in Fig. 5B and is therefore omitted. Patients are arranged three per row, with each bordered block labeled by patient identifier. Each block contains up to four stacked tracks: FFPE WGS read-depth profiles at 100-kb resolution, FFPE-CUTAC read-depth profiles at 1-Mb resolution, FFPE RNA-seq profiles inferred using CaSpER, and FF RNA-seq profiles inferred using CaSpER. FF RNA-seq tracks are shown only for patients with matched frozen tissue. In each track, the x-axis spans chromosomes 1–22, with sex chromosomes omitted, and the y-axis shows the log_2_ signal-to-baseline ratio. Gray points represent bin-level signals for WGS and FFPE-CUTAC or gene-level CaSpER signals for RNA-seq. Dashed horizontal lines mark the copy-neutral baseline at a log_2_ ratio of 1, and colored lines show segmented copy-number profiles.

**Supplementary Figure 7.**
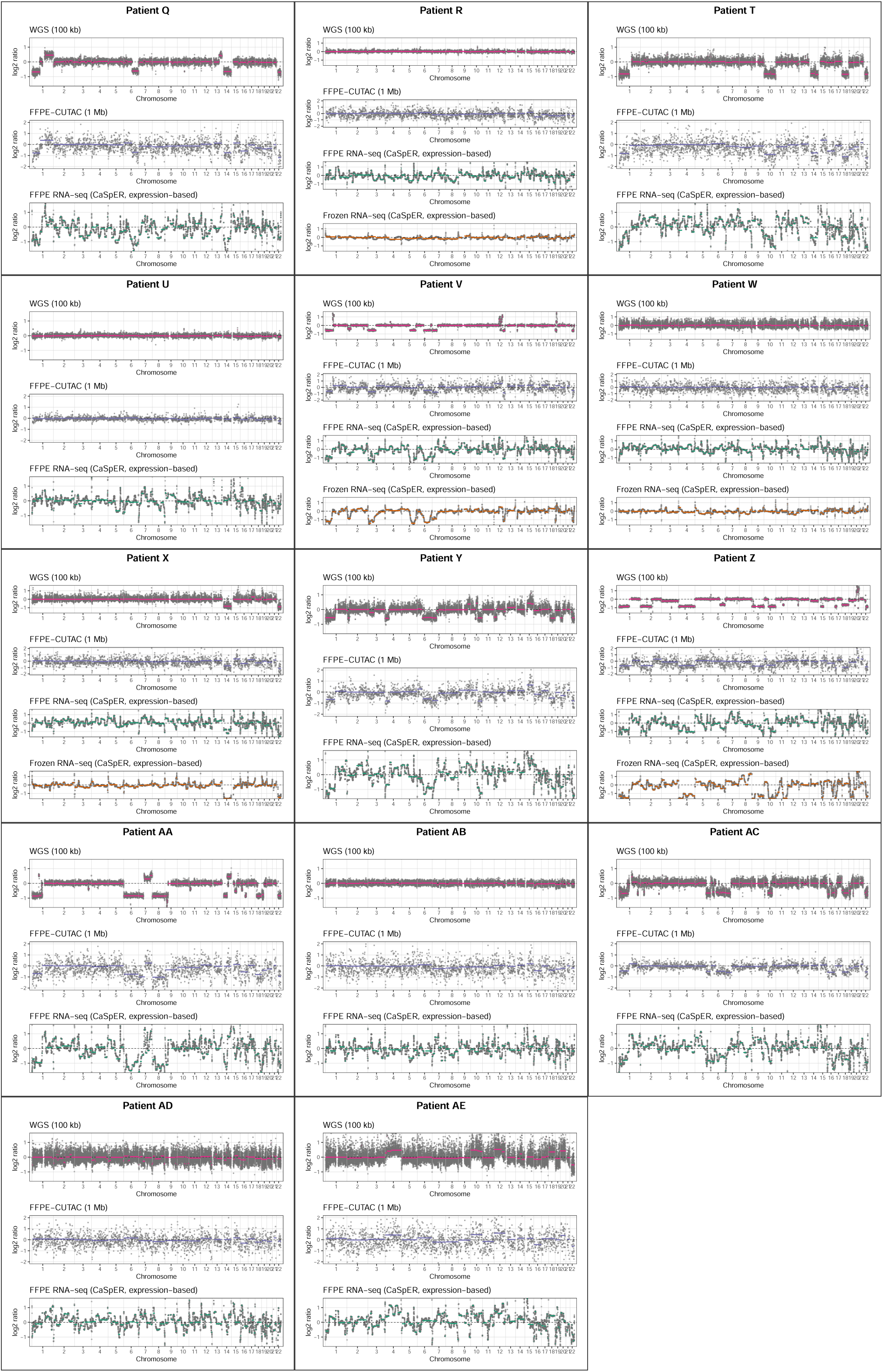
Per-patient copy-number profiles comparison across FFPE WGS, FFPE-CUTAC, and RNA-seq (continued). Continuation of the per-patient copy-number profiles shown in Supplementary Fig. 6. Profiles are shown for 14 additional meningioma patients: Q, R, T, U, V, W, X, Y, Z, AA, AB, AC, AD and AE.

**Supplementary Figure 8.**
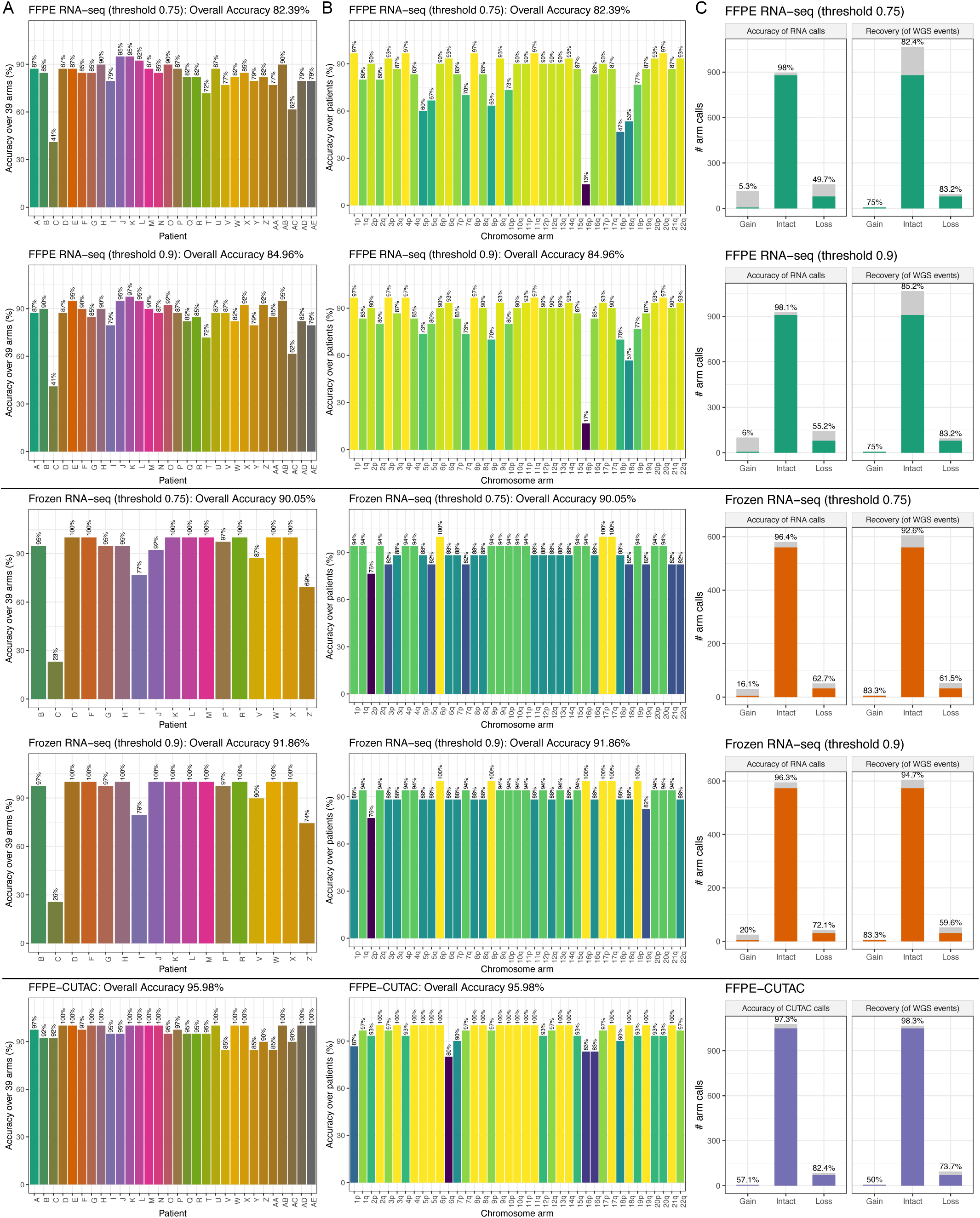
Chromosome-arm and per-patient copy-number concordance between FFPE RNA-seq, FF RNA-seq, FFPE-CUTAC and WGS. The upper and lower rows show results obtained using CaSpER^117^ thresholds of 0.75 and 0.9, respectively. **A.** Percentage of 39 autosomal chromosome arms with FFPE RNA-seq, FF RNA-seq, and FFPE-CUTAC (row 1, 2, and 3) calls concordant with WGS for each meningioma patient. Bar labels show patient-level concordance, and panel titles report overall concordance across all patient-arm calls. **B.** Percentage of patients with concordant FFPE RNA-seq, FF RNA-seq, FFPE-CUTAC and WGS calls for each chromosome arm. Bar labels show arm-level concordance. **C.** Concordance stratified by gain, intact, and loss calls. Left panels show the accuracy of FFPE RNA-seq calls, defined as the percentage of calls within each category that matched WGS. Right panels show recovery of WGS events, defined as the percentage of WGS calls within each category reproduced by FFPE RNA-seq. Colored and gray segments indicate concordant and discordant calls, respectively, and labels report the concordant percentages.

**Supplementary Figure 9.**
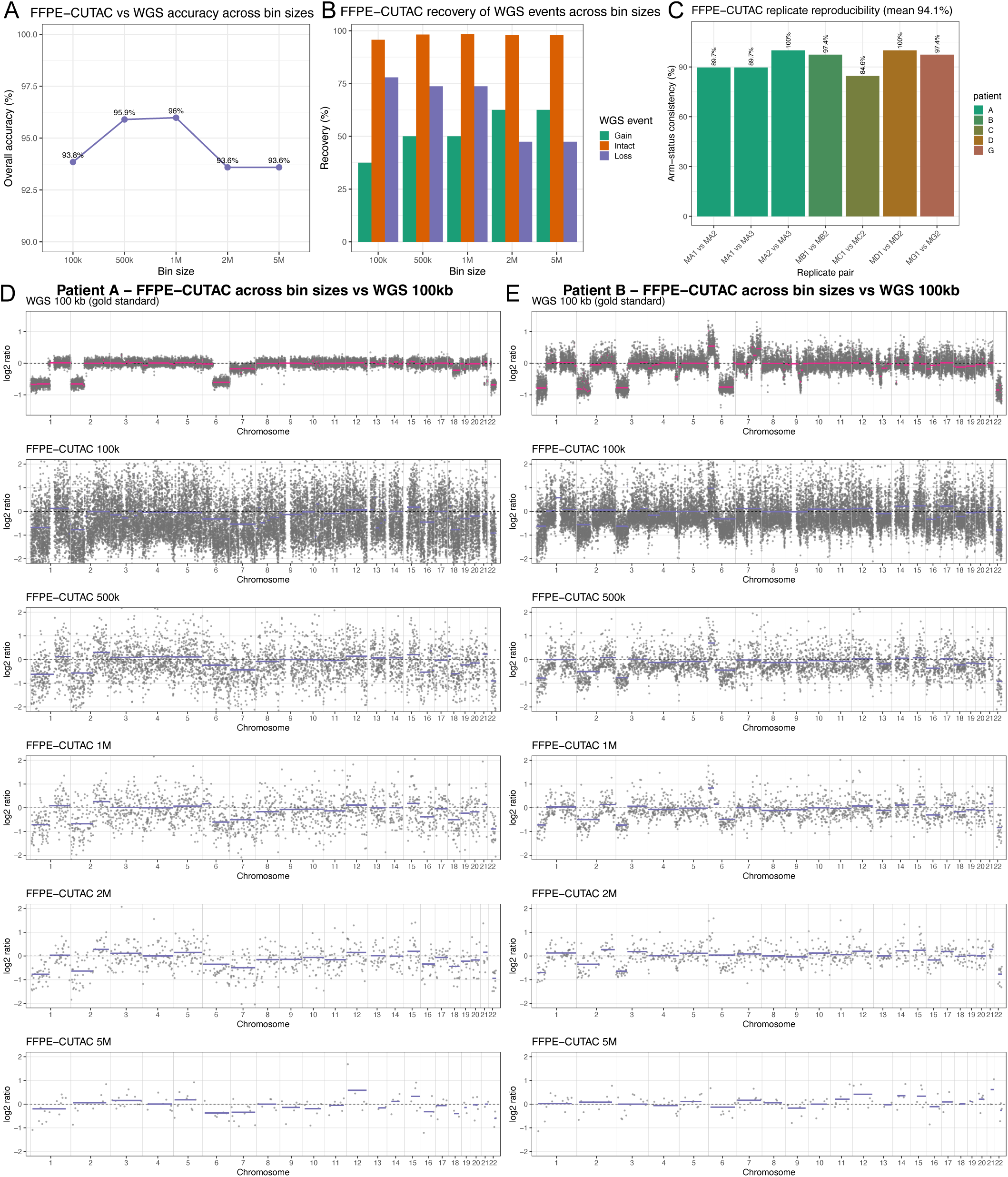
Genomic binning impact on the aneuploidy detection accuracy for FFPE-CUTAC data. **A.** Overall arm-level concordance between FFPE-CUTAC and WGS across genomic bin sizes. **B.** Recovery of WGS aneuploidy state across FFPE-CUTAC bin sizes. Bars show the percentage of WGS gain, intact and loss calls recovered by FFPE-CUTAC at each bin size. **C.** Replicates reproducibility of FFPE-CUTAC chromosome-arm calls at 1Mb resolution. **D-E.** Copy-number profiles for patient A and B across FFPE-CUTAC bin sizes. Six stacked tracks show WGS read depth at 100-kb resolution followed by FFPE-CUTAC read-depth profiles generated with 100-kb, 500-kb, 1-Mb, 2-Mb and 5-Mb bins. The x-axis shows genomic position across chromosomes 1-22. The y-axis shows the log_2_ signal-to-baseline ratio. Grey points show per-bin signals. Dashed horizontal lines mark a log_2_ ratio of 0. Colored lines show segment-level average summaries.

